# Effector target-guided engineering of an integrated domain expands the disease resistance profile of a rice NLR immune receptor

**DOI:** 10.1101/2022.06.14.496076

**Authors:** JHR Maidment, M Shimizu, S Vera, M Franceschetti, A Longya, CEM Stevenson, JC De la Concepcion, A Białas, S Kamoun, R Terauchi, MJ Banfield

## Abstract

A subset of plant intracellular NLR immune receptors detect effector proteins, secreted by phytopathogens to promote infection, through unconventional integrated domains which resemble the effector’s host targets. Direct binding of effectors to these integrated domains activates plant defences. The rice NLR receptor Pik-1 binds the *Magnaporthe oryzae* effector AVR-Pik through an integrated heavy metal-associated (HMA) domain. However, the stealthy alleles AVR-PikC and AVR-PikF avoid interaction with Pik-HMA and evade host defences. Here, we exploited knowledge of the biochemical interactions between AVR-Pik and its host target, OsHIPP19, to engineer novel Pik-1 variants that respond to AVR-PikC/F. First, we exchanged the HMA domain of Pikp-1 for OsHIPP19-HMA, demonstrating that effector targets can be incorporated into NLR receptors to provide novel recognition profiles. Second, we used the structure of OsHIPP19-HMA to guide mutagenesis of Pikp-HMA to expand its recognition profile. We demonstrate that the extended recognition profiles of engineered Pikp-1 variants correlate with effector binding in planta and in vitro, and with the gain of new contacts across the effector/HMA interface. Crucially, transgenic rice producing the engineered Pikp-1 variants were resistant to blast fungus isolates carrying AVR-PikC or AVR-PikF. These results demonstrate that effector target-guided engineering of NLR receptors can provide new-to-nature disease resistance in crops.

**Graphical abstract:** 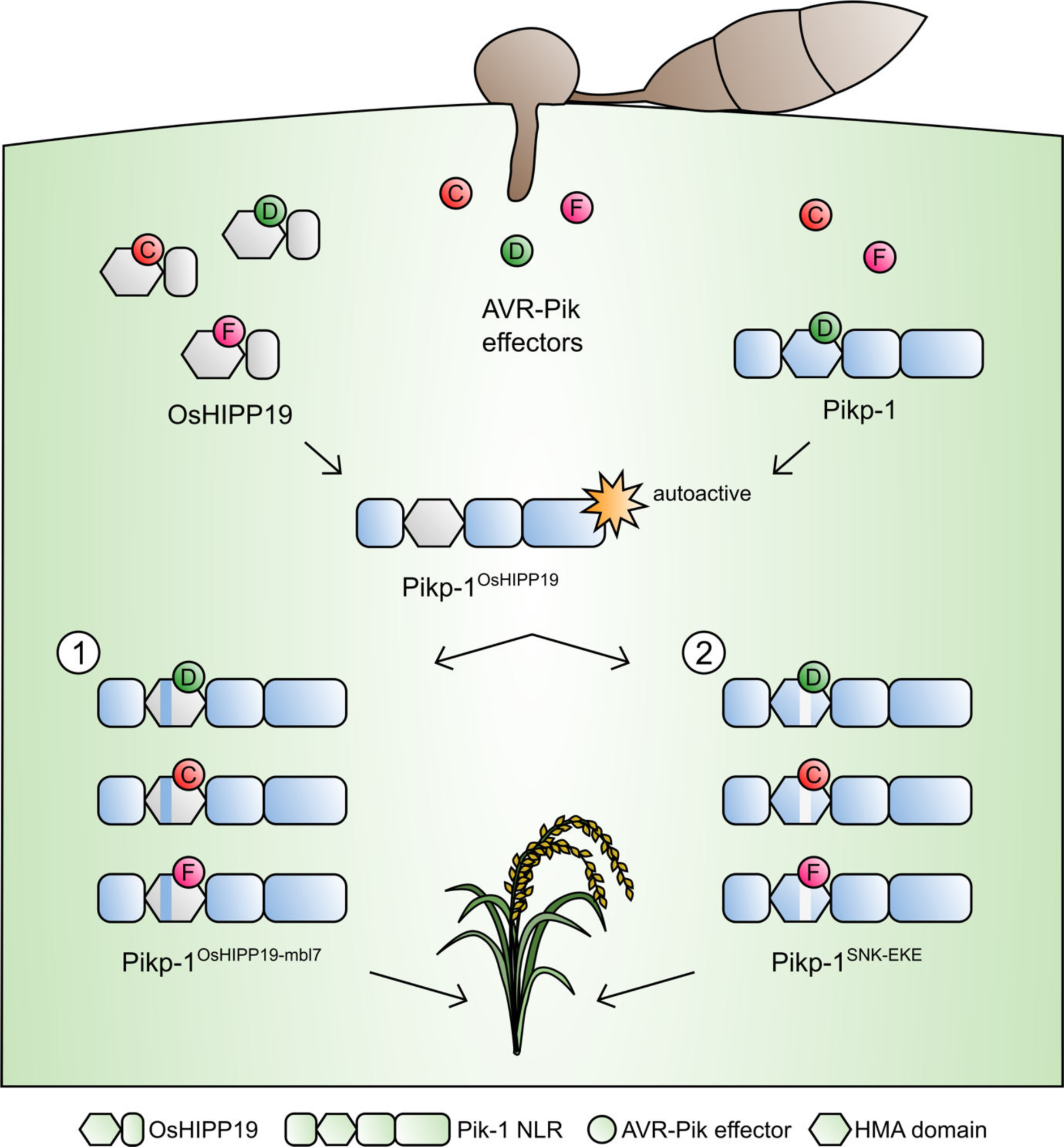

## Introduction

Intracellular <u>n</u>ucleotide-binding and leucine rich-repeat (NLR) domain-containing immune receptors are essential components of the plant innate immune system (1, 2). These receptors detect effector proteins which are delivered into host cells by invading pathogens and pests to promote virulence. NLR receptors have a modular domain architecture typically consisting of an N-terminal coiled coil (CC or CCR) or Toll/Interleukin-1 receptor (TIR) domain, a central nucleotide binding (NB-ARC) domain, and a C-terminal leucine-rich repeat (LRR) domain. In addition, some NLRs contain non-canonical domains which are integrated into the protein architecture either at the N- or C-termini or between canonical domains (3–6). These integrated domains (IDs) resemble effector virulence targets and either directly bind or are modified by effector proteins to activate NLR-mediated immune signalling (4, 7–11).

Many of the characterised resistance (R) genes used to confer disease resistance in crop breeding programmes encode NLR proteins (12). However, NLR-mediated resistance can be overcome through silencing or deletion of effectors in pathogen genomes, gain of new effectors or effector functions, or mutation to evade NLR activation (13, 14). Engineering NLRs to detect currently unrecognised effector proteins would provide new opportunities to control plant pathogens. Early attempts to engineer NLRs focused on random mutagenesis followed by gain-of-function screening, with some success in both expanding recognition profiles to new effector variants and increasing the sensitivity of the receptor (15–18). More recently, modification of the effector target PBS1, which is guarded by the NLR protein RPS5 led to successful engineering of novel recognition by this system (19–22). RPS5 is activated by cleavage of the *Arabidopsis thaliana* protein kinase PBS1 by the *Pseudomonas syringae* effector AvrPphB. By varying the PBS1 cleavage site, the RPS5/PBS1 system has been engineered to recognise proteases from different pathogens (20, 22). Using a different strategy, a protein domain targeted for degradation by the phytoplasma effector SAP05 was fused to the C-terminus of the TIR-NLR RRS1-R. RRS1-R represses the immune cell death-triggering activity of a second TIR-NLR, RPS4. While transient co-expression of the engineered RRS1-R, RPS4 and the phytoplasma effector SAP05 led to cell death in *N. tabacum*, transgenic *A. thaliana* plants were not resistant to phytoplasma carrying the SAP05 effector (23).

Integrated domains can facilitate recognition of structure- and sequence-diverse effectors which target similar host proteins. This is exemplified by the TIR-NLR pair RRS1 and RPS4, which mediate recognition of the structurally distinct effectors PopP2 from *Ralstonia solanacearum* and AvrRps4 from *Pseudomonas syringae* pv. *pisi* (10, 11, 24–27). Furthermore, the potential to replace naturally occurring integrated domains with nanobodies of defined specificity to confer disease resistance has recently been demonstrated (28).

Modification of existing integrated domains, or the incorporation of entirely new protein domains into an NLR structure could deliver new recognition specificities and extend the toolbox of resistance genes available to combat crop pathogens and pests.

The paired rice CC-NLR proteins Pik-1 and Pik-2 cooperatively activate plant defence in response to the blast pathogen effector AVR-Pik (29–31). The sensor NLR Pik-1 contains an integrated heavy metal associated (HMA) domain between the CC and NB-ARC domains (9) (figure S1a). Direct binding of AVR-Pik to the HMA domain is required to activate Pik-mediated immunity (9, 32, 33). Multiple Pik alleles have been described in different rice cultivars, with most amino acid polymorphisms located within the integrated HMA domain of Pik-1. Five Pik alleles (Pikp, Pikm, Pikh, Piks and Pik*) have been functionally characterised for their response to blast isolates carrying different AVR-Pik variants (9, 30, 32, 33) (figure S1b). To date, six AVR-Pik variants (A-F) have been described, which differ in five amino acid positions at the HMA-binding interface (30, 34, 35) (figure S1c). These polymorphisms influence binding of the effector to the integrated HMA domain of Pik-1 (9, 32, 35). Interestingly, the Asp67 and Lys78 polymorphisms of AVR-PikC and AVR-PikF, respectively, disrupt interactions between the effector and all tested integrated Pik-HMA domains (8, 33, 35). To date, none of the characterised Pik alleles can confer disease resistance to blast isolates carrying AVR-PikC or AVR-PikF (9, 30, 32, 33).

The molecular basis of interaction between AVR-Pik effectors and the integrated HMA domains of Pikp-1, Pikm-1 and Pikh-1 has been well explored (9, 32, 33). Pikp-1 is only able to recognise the AVR-PikD variant, however the introduction of two amino acid changes (Asn261Lys and Lys262Glu) extends recognition to AVR-PikE and AVR-PikA, phenocopying the recognition profile of Pikm-1 and Pikh-1 (36).

The NLR pair RGA5 and RGA4 detect the blast pathogen effectors AVR-Pia and AVR1-CO39, with activation requiring binding of the effector to an integrated HMA domain at the C-terminus of RGA5 (37–39). Crystal structures of the RGA5-HMA/AVR1-CO39 and Pik-HMA/AVR-Pik complexes were used to engineer the RGA5-HMA domain to bind AVR-PikD in addition to its cognate effectors AVR-Pia and AVR1-CO39 and deliver cell death in *Nicotiana benthamiana*, but not disease resistance in transgenic rice (40). More recently, RGA5 has been engineered to bind the non-cognate effector AVR-Pib. Transgenic rice carrying the engineered RGA5 variant was resistant to AVR-Pib-expressing *M. oryzae* strains, with resistance comparable to that displayed by the (untransformed) rice cultivar K14 which carries the Pib CC-NLR resistance gene (41). These studies demonstrate the potential for engineering integrated domains to alter the recognition profile of the NLR protein, however engineering new-to-nature effector recognition is yet to be reported.

The AVR-Pik effector targets members of the rice heavy metal associated isoprenylated plant protein (HIPP) and heavy metal associated plant protein (HPP) families through direct interaction with their HMA domain, supporting the hypothesis that NLR integrated domains are likely derived from host proteins (7, 8). In a previous study, we showed that all AVR-Pik effector variants bind to the HMA domain of OsHIPP19 with high affinity and elucidated the structural basis of this interaction by determining the crystal structure of a OsHIPP19- HMA/AVR-PikF complex (8). This shows that effector variants which are not bound by Pik-HMA domains, and escape immune recognition, retain tight binding for HMA domains of their putative host targets.

Here, we leverage our understanding of the interaction between OsHIPP19 and AVR-Pik to engineer the integrated HMA domain of Pik-1 to expand recognition to the stealthy AVR-PikC and AVR-PikF variants, enabling new-to-nature disease resistance profiles in an NLR. We use two parallel strategies to engineer recognition. First, we demonstrate that exchanging the HMA domain of Pikp-1 for that of OsHIPP19 (including additional amino acid substitutions to prevent autoactivity), gives a chimeric Pik-1 which binds AVR-Pik effectors and triggers AVR-PikC- and AVR-PikF-dependent cell death in *N. benthamiana*. Second, guided by the structure of the OsHIPP19-HMA/AVR-PikF complex, we use targeted mutagenesis of Pikp-1 to give a second engineered Pik-1 receptor capable of binding to AVR-PikC and AVR-PikF and triggering cell death in *N. benthamiana*. Finally, we show that transgenic rice expressing either of these engineered Pik-1 proteins are resistant to blast pathogen strains carrying AVR-PikC or AVR-PikF, while rice expressing wild-type Pikp is susceptible. This work highlights how a biochemical and structural understanding of the interaction between a pathogen effector and its host target can guide rational engineering of NLR proteins with novel, and new-to-nature, disease resistance profiles.

## Results

### A Pikp-1^OsHIPP19^ chimera extends binding and response to previously unrecognised AVR-Pik variants

Previously, we reported that all AVR-Pik variants, including AVR-PikC and AVR-PikF, bind to the HMA domain of OsHIPP19 with high affinity (8). The HMA domains of Pikp-1 and OsHIPP19 share 51% amino acid identity and are structurally similar; the RMSD (as calculated in Coot using secondary structure matching) between Pikp-HMA (PDB 6G10) and OsHIPP19-HMA (PDB 7B1I) is 0.97Å across 71 amino acids. We hypothesised that exchanging the HMA domain of Pikp-1 for the HMA domain of OsHIPP19 would result in an NLR capable of binding and responding to AVR-PikC and AVR-PikF.

For this exchange, amino acids 188-263 (inclusive) of Pikp-1 were replaced with amino acids 2-77 of OsHIPP19 to give the chimeric NLR protein Pikp-1^OsHIPP19^ (figure S2a). To test whether Pikp-1^OsHIPP19^ could associate with AVR-Pik effector variants in planta, we performed co-immunoprecipitation experiments in *N. benthamiana*. Each of the myc-AVR-Pik variants (an N-terminal myc tag was used for effectors in all experiments in *N. benthamiana*) was transiently co-expressed with Pikp-1^OsHIPP19^-HF (C-terminal 6xHis/3xFLAG tag, used for all Pikp-1 constructs expressed in *N. benthamiana*) by agroinfiltration. Pikp-2 was not included to prevent the onset of cell death, which reduces protein levels in the plant cell extract, and previous work has shown that AVR-Pik can associate with Pik-1 in the absence of Pik-2 (32). Following immunoprecipitation with anti-FLAG beads to enrich for Pikp-1^OsHIPP19^, western blot analysis showed that all AVR-Pik variants co-precipitated with Pikp-1^OsHIPP19^ (figure 1a). As a control, and consistent with previous studies, AVR-PikD associated with Pikp-1, while AVR-PikC did not.

**Figure 1.**
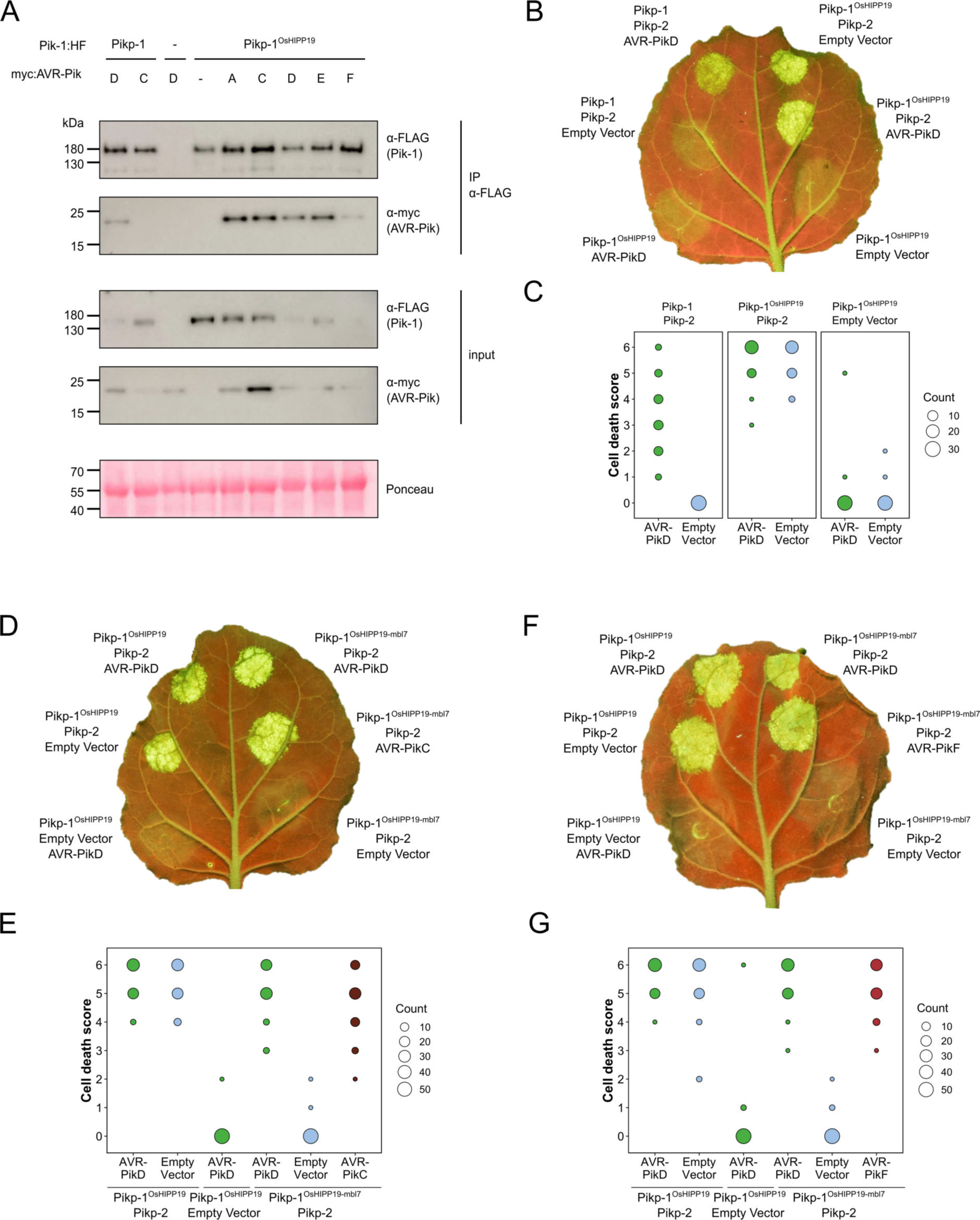
The Pikp-1^OsHIPP19-mbl7^ chimera expands binding and response to previously unrecognised AVR-Pik effector variants. (**A**) Western blots following co-immunoprecipitation revealing that the Pikp-1^OsHIPP19^ chimera associates with all AVR-Pik effector variants in *N. benthamiana*. Plant cell lysates were probed for the expression of Pikp- 1/Pikp-1^OsHIPP19^ and AVR-Pik effector variants using anti-FLAG and anti-Myc antiserum, respectively. Total protein extracts were visualised by Ponceau Staining. (**B)** Representative leaf image showing the Pikp-1^OsHIPP19^ chimera is autoactive in *N. benthamiana*. NLR-mediated responses appear as autofluorescence imaged under UV light. Pikp-mediated response to AVR-PikD (positive control, top left), Pikp-1^OsHIPP19^/Pikp-2 without effector shows autoactivity (top right), Pikp-1^OsHIPP19^/Pikp-2 response remains in the presence of AVR-PikD (middle right). Other leaf positions represent relevant negative controls. (**C**) Pikp-mediated response scoring represented as dot plots to summarise 30 repeats of the experiment shown in (**B**) across three independent experiments (Materials and Methods, figure S3). Fluorescence intensity is scored as previously described (9, 32). (**D**) The Pikp-1^OsHIPP19-mbl7^ chimera does not display autoactive cell death in *N. benthamiana* (bottom right), as seen for Pikp-1^OsHIPP19^ (middle left), but retains response to AVR-PikD and expands Pikp-mediated response to AVR-PikC (middle right). (**E**) Pikp-mediated response scoring represented as dot plots to summarise 60 repeats of the experiment shown in (**D**) across three independent experiments (Materials and Methods, figure S3). (**F**) As for (D), but showing the expanded Pikp-mediated response to AVR-PikF. (**G**) As for (E) but for 60 repeats of the experiment in (F) across three independent experiments (Materials and Methods, figure S3).

We then transiently co-expressed epitope-tagged Pik-1, Pikp-2 and AVR-Pik in *N. benthamiana* using cell death as a proxy for immune activation (9, 32, 33, 35). We found that when co-expressed with Pikp-2-HA (C-terminal hemagglutinin tag, used for all Pikp-2 constructs expressed in *N. benthamiana*), Pikp-1^OsHIPP19^ is autoactive and triggers spontaneous cell death in the absence of the effector (figure 1b, 1c, S3). This autoactivity requires an intact P-loop and MHD motif in Pikp-2, as cell death is abolished when Pikp- 1^OsHIPP19^ is transiently co-expressed with either Pikp-2^K217R^ or Pikp-2^D559V^ (figure S4a, S4b, S5). Cell death was reduced, but not abolished, when Pikp-1^OsHIPP19^ with a Lys296Arg mutation in the P-loop motif (Pikp-1^OsHIPP19_K296R^) was transiently co-expressed with Pikp-2 (figure S4a, S4b, S5). Western blot analysis indicated that all fusion proteins were produced (figure S4c, S6a).

A previous study showed that autoactivity following HMA domain exchange could be abolished by reverting the degenerate metal-binding motif of the HMA domain (“MxCxxC”) to the corresponding amino acids in Pikp-1 (42). Based on this observation, we exchanged seven amino acids (encompassing the entire MxCxxC motif) in the β1-α1 loop of the Pikp-1^OsHIPP19^ chimera for the corresponding amino acids in Pikp-1 (figure S2b, S7). The resulting chimera, Pikp-1^OsHIPP19-mbl7^ hereafter (mbl7 refers to 7 amino acids in the “<u>m</u>etal-<u>b</u>inding loop”), was not autoactive, and did not trigger spontaneous cell death in the absence of the effector. Crucially, Pikp-1^OsHIPP19-mbl7^ retained the ability to trigger cell death in *N. benthamiana* when co-expressed with Pikp-2 and AVR-PikD. Further, Pikp-1^OsHIPP19-mbl7^ also triggered cell death in *N. benthamiana* when co-expressed with Pikp-2 and AVR-PikC or AVR-PikF (figure 1d-g, S3b, S3c, S6b, S6c, S8). We confirmed that Pikp-1^OsHIPP19-mbl7^ retains binding to all AVR-Pik variants by co-immunoprecipitation in *N. benthamiana* as for Pikp-1^OsHIPP19^ (figure S9). Western blot analysis showed that all proteins for the cell death assays were produced in leaf tissue (figure S6b, S6c).

### Structure-guided mutagenesis of Pikp-1 extends response to previously unrecognised AVR-Pik variants

Alongside the HMA-domain exchange strategy, we also used a structure-guided approach to target point mutations in Pikp-HMA that could extend the effector recognition profile of Pikp without triggering autoimmunity.

The interaction surfaces between integrated Pik-HMA domains and AVR-Pik effectors are well-characterised, with crystal structures revealing three predominant interfaces (termed 1-3) between the proteins (9, 32, 33, 36). These interfaces are also observed in the structure of the HMA domain of OsHIPP19 in complex with AVR-PikF (PDB accession code 7B1I (8)). The Asp67 and Lys78 polymorphisms that distinguish AVR-PikC and AVR-PikF from AVR-PikE and AVR-PikA, respectively, are located at interface 2. In the crystal structure of Pikh-HMA/AVR-PikC, the sidechain of AVR-PikC^Asp67^ extends towards a loop in the HMA domain containing Pikh-HMA^Asp224^. This loop is shifted away from the effector, likely due to steric clash and/or repulsion by the two Asp sidechains, and intermolecular hydrogen bonds between Pikh-HMA^Asp224^ and AVR-PikC^Arg64^ are disrupted. We hypothesised that compensatory mutations at interface 2 could mitigate against the disruption caused by AVR-PikC^Asp67^. Therefore, we introduced Asp224Ala and Asp224Lys mutations in the Pikp-1^NK-KE^ background and tested these constructs in cell death assays in *N. benthamiana*. Neither mutation extended the Pikp-1^NK-KE^-mediated cell death response to AVR-PikC, and both mutations reduced the extent of the response to AVR-PikD (figure S10, S11).

Pikh-1 and Pikp-1^NK-KE^ differ from Pikp-1 by one and two amino acids, respectively, at interface 3. These amino acid differences are sufficient to extend binding and cell death response to AVR-PikE and AVR-PikA, even though the residues that distinguish these variants from AVR-PikD are located at interface 2. We therefore predicted that we could engineer a modified Pik- 1 that interacts with AVR-PikC/AVR-PikF by mutating other interfaces in the HMA domain to compensate for disruption at the site of the polymorphic residue. The crystal structure of the OsHIPP19-HMA/AVR-PikF complex revealed additional hydrogen bond interactions at interface 3 relative to integrated HMAs in complex with AVR-Pik variants (8). The side chain of OsHIPP19^Glu72^ was particularly striking. The corresponding residue in all described Pik-1 HMA domains is serine, and while the hydroxyl group of the serine side chain only forms an intramolecular hydrogen bond within the HMA domain, the bulkier OsHIPP19^Glu72^ side chain extends across the interface and forms a direct hydrogen bond with the effector (figure 2a). We therefore introduced a Ser258Glu mutation in the Pikp-1^NK-KE^ background to give the triple mutant Pikp-1^SNK-EKE^ and tested the ability of this protein to respond to AVR-Pik variants in *N. benthamiana* cell death assays. Firstly, we confirmed that Pikp-1^SNK-EKE^ was not autoactive, evidenced by a lack of cell death following co-expression with Pikp-2 only (in the absence of effectors). Next, we established that Pikp-1^SNK-EKE^ remained functional and caused cell death on co-expression with Pikp-2 and AVR-PikD. Crucially, transient expression of Pikp-1^SNK-EKE^, but not Pikp-1^NK-KE^, with Pikp-2 and either AVR-PikC or AVR-PikF, triggered cell death suggestive of new effector response specificities (figure 2b, 2c, 2d, 2e, S12, S13). Western blot analysis indicated that all fusion proteins were produced (figure S14).

**Figure 2.**
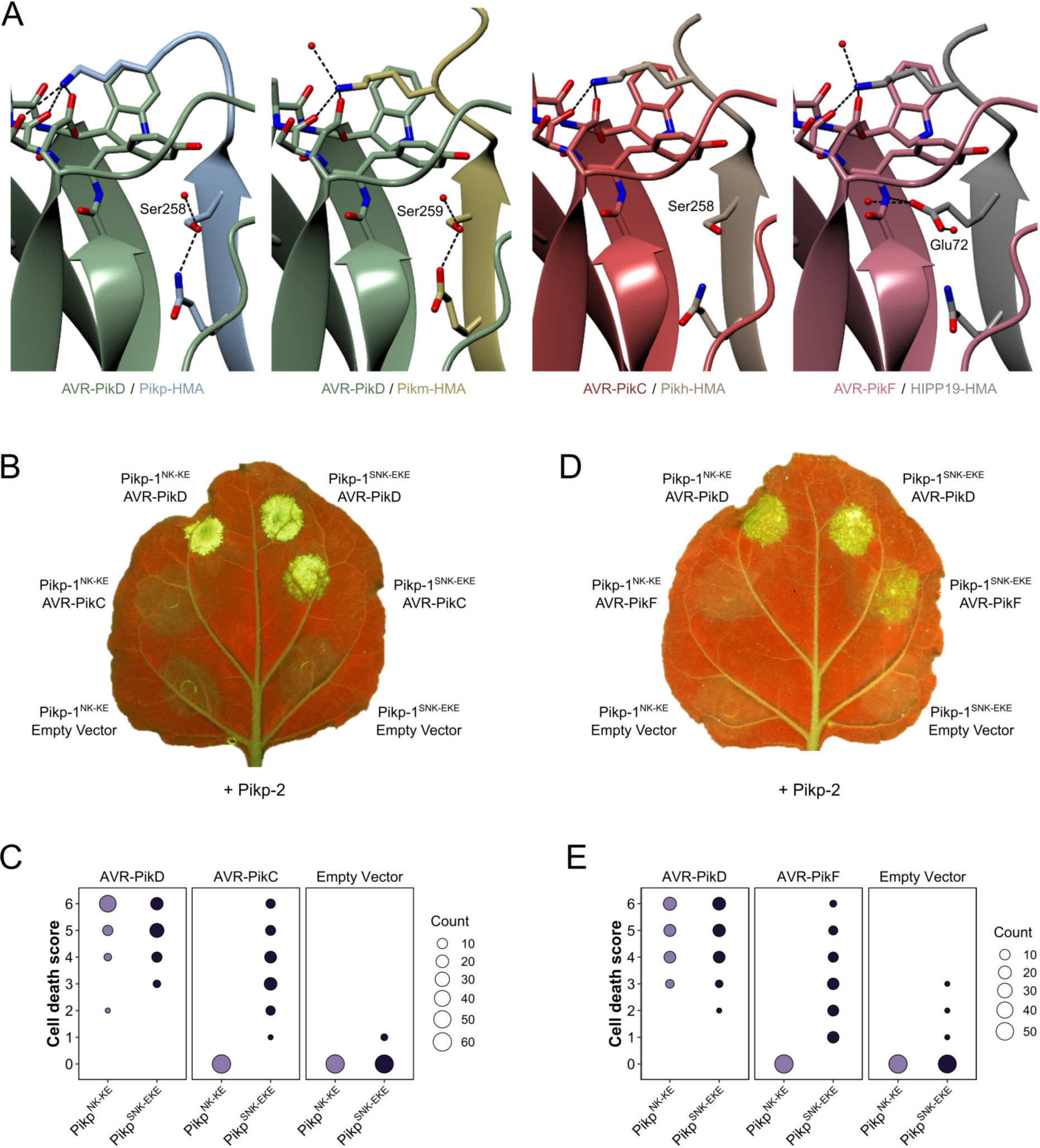
Structure-guided mutagenesis of Pikp-1 expands response to previously unrecognised AVR-Pik effector variants. (**A**) Comparison of the crystal structures of AVR- Pik effector variants in complex with Pik-HMA domains (PDB entries 6G10, 6FU9 and 7A8X) and AVR-PikF in complex OsHIPP19 (PDB entry 7B1I) suggests addition of an S258E mutation to the NK-KE mutations described previously (36) could introduce new contacts across the protein:protein interface. Protein structures are represented as ribbons with relevant side chains displayed as cylinders. Dashed lines indicate hydrogen bonds. Relevant water molecules are represented as red spheres. (**B**) The Pikp^SNK-EKE^ mutant gains response to AVR-PikC (right, middle) where no response is observed for Pikp^NK-KE^ (left, middle). Further, the Pikp^SNK-EKE^ mutant is not autoactive (right, bottom) and retains response to AVR-PikD (right, top). All infiltration spots contain Pikp-2. (**C**) Pikp-mediated response scoring represented as dot plots to summarise 60 repeats of the experiment shown in (**B**) across three independent experiments (Materials and Methods, figure S12). (**D**) and (**E**) as described for and (**C**) but with AVR-PikF and 57 repeats across three independent experiments.

We tested whether the Ser258Glu mutation alone was sufficient to extend the cell death response to AVR-PikC or AVR-PikF using the cell death assay. When Pikp-1^S258E^ was co- infiltrated with Pikp-2 and either AVR-PikC or AVR-PikF no cell death was observed (figure S15, S16), demonstrating that the triple mutation is necessary for response to these effectors.

### The Ser258Glu mutation extends binding of Pikp-HMA^NK-KE^ to AVR-PikC and AVR-PikF in vitro

The extent of the Pik/AVR-Pik-dependent cell death response in *N. benthamiana* largely correlates with binding affinity in vitro and in planta (9, 32, 33, 36). To test whether the Pikp- 1^SNK-EKE^ response to AVR-PikC or AVR-PikF in *N. benthamiana* correlates with increased binding to the modified HMA domain, we first used surface plasmon resonance (SPR) with purified proteins. Pik-HMA domains and AVR-Pik effectors were purified from *E. coli* cultures using established protocols for production of these proteins (8, 9, 32, 33, 36). For SPR, AVR-Pik effector variants were immobilised on a Ni^2+^-NTA sensor chip via a C-terminal 6xHis tag.

Pikp-HMA, Pikp^NK-KE^-HMA or Pikp^SNK-EKE^-HMA was flowed over the surface of the chip at three different concentrations (4nM, 40nM and 100nM). The binding (Robs, measured in response units, RU) was recorded and expressed as a percentage of the maximum theoretical responses (%Rmax), assuming a 2:1 HMA:effector interaction model (9). Consistent with previous studies, Pikp-HMA did not bind AVR-PikC (nor AVR-PikF), and weak binding was observed for Pikp^NK-KE^-HMA to AVR-PikC (and also to AVR-PikF). By contrast, Pikp^SNK-EKE^- HMA bound to both AVR-PikC and AVR-PikF with higher apparent affinity (larger %Rmax) than Pikp^NK-KE^-HMA (figure 3a, 3b). This result was consistent across the three concentrations investigated, though binding (and %Rmax) was low for all three HMA domains at 4nM (figure S17).

**Figure 3.**
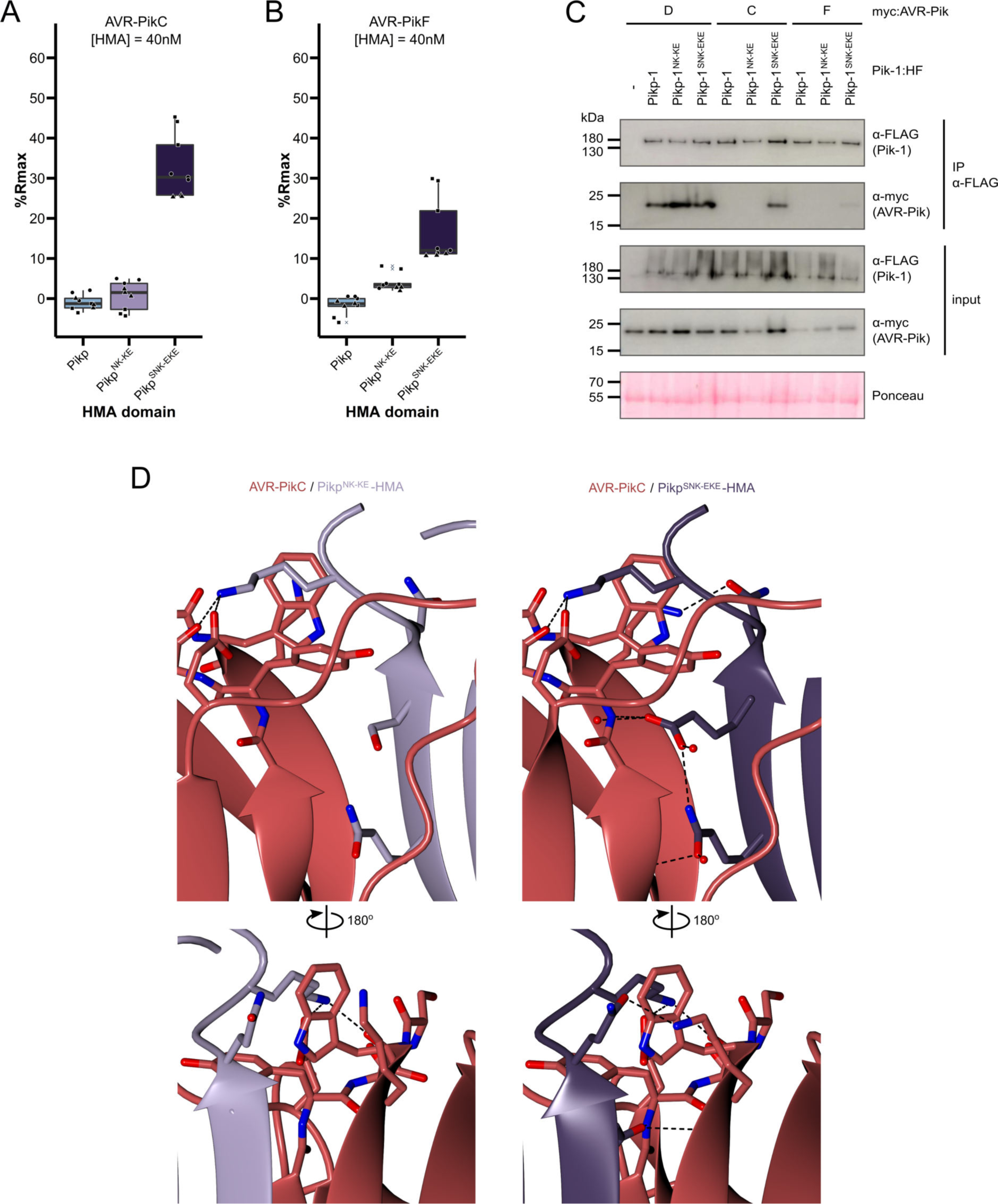
The SNK-EKE triple mutation extends Pikp-1 binding to AVR-PikC and AVR- PikF in vitro and in planta by facilitating new contacts across the protein:protein interface. Boxplots showing the %Rmax observed for the interactions between AVR-PikC (**A**) or AVR-PikF (**B**), both at 40nM injection concentration, and each of Pikp-HMA, Pikp-HMA^NK- KE^ and Pikp-HMA^SNK-EKE^. %Rmax is the percentage of the theoretical maximum response, assuming a 2:1 binding model (as previously observed for Pikp-HMA proteins).The center line of the box represents the median and the box limits are the upper and lower quartiles. The whiskers extend to the smallest value within Q1 − 1.5Å∼ the interquartile range (IQR) and the largest value within Q3 + 1.5Å∼ IQR. Individual data points are represented as black shapes. The experiment was repeated three times, with each experiment consisting of three technical replicates. Data for 4nM and 100nM effector injection concentrations are shown in figure S17. Western blots following co-immunoprecipitation show that the Pikp-1^SNK-EKE^ chimera binds to tested AVR-Pik effector variants in *N. benthamiana*. Plant cell lysates were probed for the expression of Pikp-1/Pikp-1^NK-KE^/ Pikp-1^SNK-EKE^ and AVR-Pik effector variants using anti-FLAG and anti-Myc antiserum, respectively. Total protein extracts were visualised by Ponceau Staining. (**D**) The crystal structure of the Pikp-HMA^SNK-EKE^/AVR-PikC complex (PDB entry 7QPX) reveals additional hydrogen bonds at the protein:protein interfaces compared to Pikp- HMA^NK-KE^/AVR-PikC (PDB entry 7A8W). Protein structures are represented as ribbons with relevant side chains displayed as cylinders. Dashed lines indicate hydrogen bonds. Relevant water molecules are represented as red spheres.

### The Ser258Glu mutation extends binding of Pikp-HMA^NK-KE^ to AVR-PikC and AVR-PikF in planta

Next, we determined whether the Ser258Glu mutation also extends binding to AVR-PikC and AVR-PikF in the full length NLR in planta using co-immunoprecipitation. Full-length Pikp-1, Pikp-1^NK-KE^ and Pikp-1^SNK-EKE^ were each co-expressed with either AVR-PikD, AVR-PikC, or AVR-PikF in *N. benthamiana*. As before, Pikp-2 was not included in the co-immunoprecipitation assays to prevent the onset of cell death. We found that AVR-PikD, AVR-PikC or AVR-PikF co-immunoprecipitated with Pikp-1^SNK-EKE^ (figure 3c); while the band corresponding to AVR-PikF was faint, this can be attributed to lower levels of AVR-PikF in the input. As previously observed (36), AVR-PikD, but not AVR-PikC or AVR-PikF co-immunoprecipitated with Pikp-1 and Pikp-1^NK-KE^ (figure 3c). Taken together, the results from in vitro and in planta assays indicate that the Ser258Glu mutation increases the binding of Pikp-1^NK-KE^ for AVR- PikC and AVR-PikF to a sufficient level to trigger cell death in planta.

### Crystal structures of the Pikp-HMA^SNK-EKE^/AVR-PikC and Pikp-HMA^SNK-EKE^/AVR-PikF complexes reveal new contacts across the binding interface

To confirm that the side chain of Glu258 in Pikp-1^SNK-EKE^ forms a new hydrogen bond across the interface (as observed for Glu72 in the OsHIPP19/AVR-PikF complex), we determined the crystal structures of the Pikp-HMA^SNK-EKE^/AVR-PikC and Pikp-HMA^SNK-EKE^/AVR-PikF complexes. For comparison, we also determined the crystal structure of Pikp-HMA^NK-KE^/AVR- PikC. These were produced by co-expression in *E. coli* and purified to homogeneity using established methods for purification of HMA domain/AVR-Pik complexes (8, 9, 32, 36). Crystals were obtained in several conditions in the Morpheus® screen (Molecular Dimensions), and X-ray diffraction data were collected at the Diamond Light Source (Oxford, UK) to a resolution of 2.15 Å (Pikp-HMA^NK-KE^/AVR-PikC), 2.05 Å (Pikp-HMA^SNK-EKE^/AVR-PikC) and 2.2 Å (Pikp-HMA^SNK-EKE^/AVR-PikF). These structures were solved by molecular replacement and refined/validated using standard protocols (see Materials and Methods). Data collection, processing and refinement statistics are shown in table S1. The final refined models have been deposited at the PDB with accession codes 7A8W, 7QPX and 7QZD.

The global structure of the complexes are essentially identical to each other and to the previously determined Pik-HMA/AVR-Pik crystal structures (figure S18, S19, S20, table S1). The RMSDs, as calculated in COOT with secondary structure matching, between Pikp-HMA^NK- KE^/AVR-PikC and Pikp-HMA^SNK-EKE^/AVR-PikC or Pikp-HMA^SNK-EKE^/AVR-PikF are 0.38 Å using 154 residues and 0.60 Å using 155 residues, respectively. Interface analysis performed with qtPISA (43) identified 15 hydrogen bonds and 9 salt bridges between Pikp-HMA^SNK-EKE^ and AVR-PikC, and 16 hydrogen bonds and 11 salt bridges between Pikp-HMA^SNK-EKE^ and AVR- PikF, compared to the 12 hydrogen bonds and 8 salt bridges mediating the interaction between Pikp-HMA^NK- KE^ and AVR-PikC (table S2). Inspection of the structures revealed that the side chain of Glu258 does indeed extend across the interface, forming direct hydrogen bonds with the backbone of the effectors (figure 3d). This single mutation also supports additional hydrogen bonds at the interface between AVR-PikC (or AVR-PikF) and residues comprising β4 of the HMA domain (figure 3d). These differences at interface 3 likely explain the increased binding affinity of Pikp-HMA^SNK-EKE^ for AVR-PikC/AVR-PikF relative to Pikp- _HMA_NK-KE.

### Rice plants expressing Pikp^OsHIPP19-mbl7^ or Pikp-1^SNK-EKE^ are resistant to *M. oryzae* expressing AVR-PikC or AVR-PikF

To determine whether the engineered Pik NLRs Pikp-1^OsHIPP19-mbl7^ and Pikp-1^SNK-EKE^ could mediate resistance to *M. oryzae* Sasa2 isolates expressing either AVR-PikC or AVR-PikF, we generated transgenic rice (*Oryza sativa* cv. Nipponbare, *pikp-*) expressing either wild-type *Pikp-1*, *Pikp-1^OsHIPP19-mbl7^* or *Pikp-1^SNK-EKE^*, with *Pikp-2*. Expression of both *Pikp-1* (or engineered *Pikp-1* variants) and *Pikp-2* were under the control of the constitutive CaMV 35S promoter and confirmed by RT-PCR (figure S24). Rice leaf blade punch inoculation assays were performed to determine resistance to *M. oryzae* (Sasa2) transformants carrying *AVR-Pik* alleles (*AVR-PikC, -PikD* or -*PikF*) or *AVR-Pii* (negative control, AVR-Pii is not recognised by Pikp) in T1 progenies derived from one *Pikp-1/Pikp-2*, six *Pikp-1^OsHIPP19-mbl7^/Pikp-2* and five *Pikp-1^SNK-EKE^/Pikp-2* independent transgenic T0 lines (figure 4, S21, S22, S23). The T1 transgenic rice lines expressing *Pikp-1/Pikp-2* showed resistance to Sasa2 transformed with AVR-PikD, but not to the other transformants (Figure 4AB, figure S13A, figure S14). In contrast, the T1 transgenic rice lines expressing either Pikp-1^OsHIPP19-mbl7^/Pikp-2 or Pikp-1^SNK- EKE^/Pikp-2 were resistant to Sasa2 transformed with either AVR-PikC or AVR-PikF, as well as to Sasa2 transformed with AVR-PikD (figure 4, S21, S22, S23).

**Figure 4.**
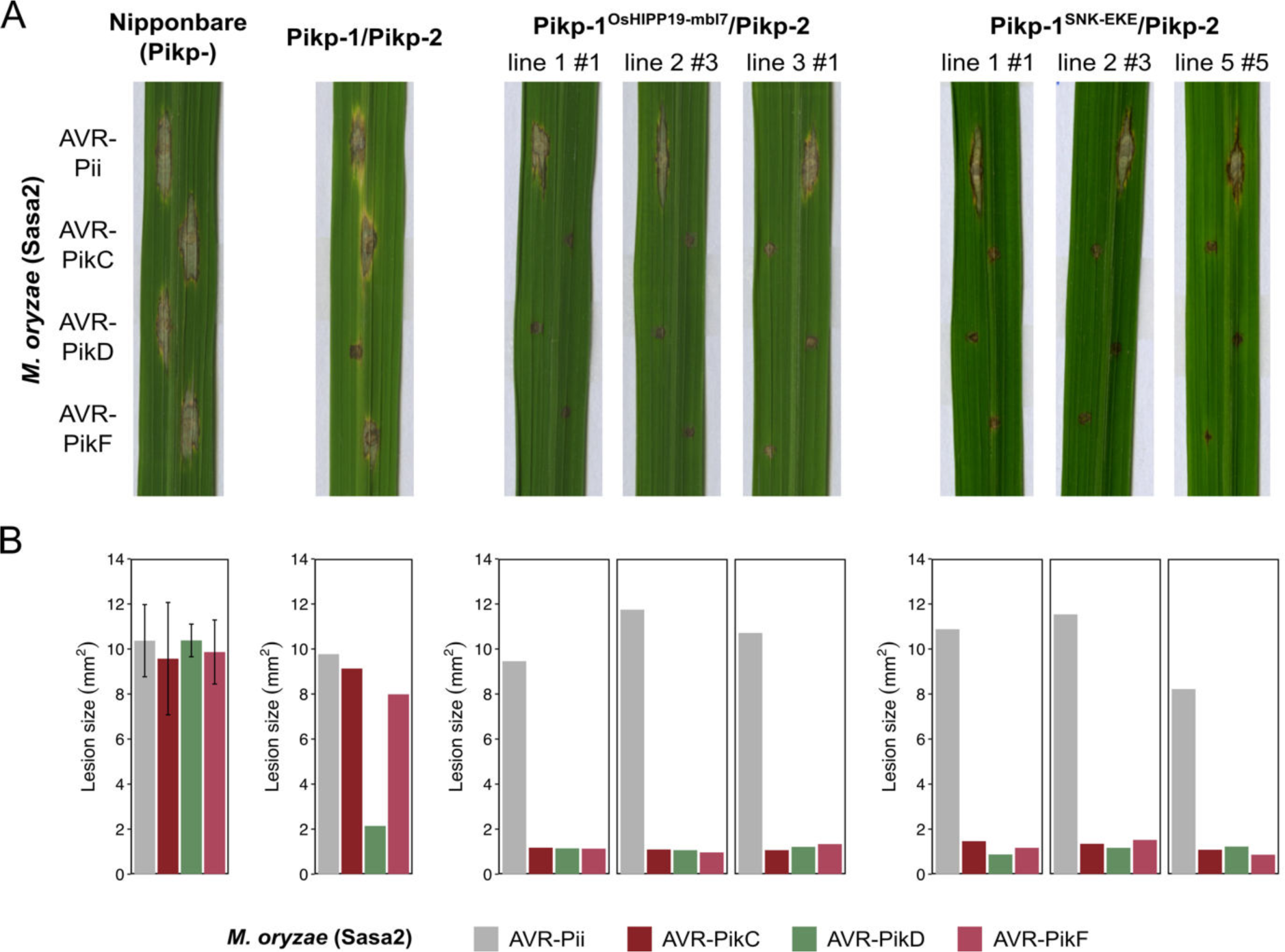
Transgenic rice plants carrying the *Pikp-1^OsHIPP19mbl7^* chimera or the *Pikp-1^SNK- EKE^* mutation show extended resistance to *Magnaporthe oryzae* carrying *AVR-PikC* or *AVR-PikF* compared to Pikp-1 wild-type. (A) Example leaves from pathogenicity assays of wild-type *O. sativa* cv. Nipponbare and three transgenic lines of *O. sativa* cv. Nipponbare expressing *Pikp-1/Pikp-2*, *Pikp-1^OsHIPP19mbl7^/Pikp-2* or *Pikp-1^SNK-EKE^/Pikp-2* challenged with *M. oryzae* Sasa2 transformed with *AVR-Pii*, *AVR-PikC*, *AVR-PikD* or *AVR-PikF*. The T1 generation seedlings were used for the inoculation test. Wild-type *O. sativa* cv. Nipponbare (recipient) is susceptible to all *M. oryzae* Sasa2 transformants (left), while the *Pikp-1/Pikp-2* transformant is only resistance to *M. oryzae* Sasa2 transformed with *AVR-PikD* (no development of disease lesions). The *Pikp-1^OsHIPP19mbl7^/Pikp-2* or *Pikp-1^SNK-EKE^/Pikp-2* plants show resistance to *M. oryzae* Sasa2 transformed with *AVR-PikC, AVR-PikD* or *AVR-PikF* but not *AVR-Pii.* (**B**) Disease lesion sizes (determined using ImageJ) represented as bar charts. For Nipponbare, the chart shows the mean average lesion size for 5 leaves, with error bars showing SE. All other chart are for the specific leaves shown and are not averaged across repeats to account for different genetic backgrounds. Repeat experiments in different lines are shown in figures S21-23. RT-PCR confirming expression of transgenes is shown in figure S24.

## Discussion

Plant diseases cause significant crop losses and constrain global food production. To develop disease resistant crops, breeding programmes exploit resistance genes present in wild germplasm that can be introgressed into elite cultivars. While recent advances have accelerated efforts to identify and clone resistance genes (44–47), conventional breeding approaches are constrained by the recognition profiles of resistance genes present in wild germplasm. Rational engineering of NLR immune receptors has the potential to yield novel disease resistance traits and expand the repertoire of resistance genes available to combat plant pathogens. It also offers the potential to restore disease resistance that has been overcome by pathogens and accelerate responses to dynamic changes in pathogen effector populations. Here, we took two approaches to engineer the integrated HMA domain of the NLR protein Pik-1 to deliver new-to-nature effector recognition profiles.

The stealthy effector variants AVR-PikC and AVR-PikF do not interact with the integrated HMA domains of any Pik alleles characterised to date with sufficiently high affinity to activate defence. By contrast, as a putative virulence target of AVR-Pik, OsHIPP19 is bound by all effector variants, including AVR-PikC and AVR-PikF with high affinity (8, 48). Using this knowledge, and the relationship between OsHIPP19 and integrated Pik-HMA domains, we engineered two Pik-1 variants, Pikp-1^OsHIPP19-mbl7^ and Pikp-1^SNK-EKE^. These engineered Pik-1 proteins bound AVR-PikC and AVR-PikF, activated cell death in *N. benthamiana*, and conferred blast resistance in rice. Engineering an NLR integrated domain to resemble an effector target reduces the likelihood of the effector mutating to evade immune detection while retaining host target binding. Therefore, this approach may represent a route to more durable disease resistance, particularly in the case of effectors whose function is essential for pathogen virulence.

Despite the structural similarity of the OsHIPP19 and Pikp-1 HMA domains, the Pikp-1^OsHIPP19^ chimera triggered effector-independent cell death in *N. benthamiana* when expressed with Pikp-2. This autoactivity required intact P-loop and MHD motifs in Pikp-2, as previously observed for the effector-dependent response of Pikp-1/Pikp-2 to AVR-PikD (31). Interestingly, mutating the conserved lysine in the P-loop of Pikp-1^OsHIPP19^ to arginine partially attenuated the cell death response. Mutating the P-loop of Pikp-1 has previously been shown to abolish the effector-dependent cell death response; however maintenance of effector-independent cell death, albeit at a reduced level, by Pikp-1^OsHIPP19_K296R^, suggests that an intact P-loop in Pikp-1 is not essential for Pik-mediated signalling. Based on a previously published approach to remove autoactivation when incorporating different HMA domains into Pik-1 (42), we reverted seven amino acids in the β1-α1 loop of Pikp-1^OsHIPP19^ to those found in Pikp-1, giving the modified chimera Pikp-1^OsHIPP19-mbl7^. The β1-α1 loop contains the classical MxCxxC metal-binding motif which is characteristic of HMA domains and is degenerate in both Pikp-1 (MEGNNC) and OsHIPP19 (MPCEKS). We speculate that this loop is involved in intra- or inter- molecular interactions of Pik-1/Pik-2 which support an inactive state in the absence of effector binding, which are disturbed in the Pikp-1^OsHIPP19^ chimera but restored in Pikp-1^OsHIPP19-mbl7^. These results highlight potential challenges of incorporating domains which have not co-evolved with other domains in the receptor, but also shows the potential for overcoming autoactivation to deliver functional NLRs. A recent study demonstrated that the Pik-1 chassis can accommodate nanobodies to GFP and mCherry, and these mediated reduced viral loads of Potato Virus X (PVX) expressing these antigens in *N. benthamiana* (28). This further demonstrates the potential of the Pik-1 NLR as a versatile system for engineering disease resistance through domain exchange.

Based on the OsHIPP19/AVR-PikF complex (8), we incorporated a Ser258Glu point mutation in the Pikp-1^NK-KE^ background, generating a Pikp-HMA triple mutant, Pikp-1^SNK-EKE^. By determining the crystal structures of the Pikp-HMA^SNK-EKE^/AVR-PikC and Pikp-HMA^SNK- EKE^/AVR-PikF complexes, we confirmed the formation of new contacts across the HMA/effector interface that likely account for the expanded recognition profile to AVR-PikC and AVR-PikF. In addition to a new hydrogen bond between the side chain of Pikp- 1^SNK- EKE_Glu258^ and the backbone of AVR-PikC, we observed two additional intermolecular hydrogen bonds formed between other amino acids at the interface. This extended hydrogen bonding is facilitated by a shift in β4 of the HMA domain towards the effector. Together with previous studies in the Pik-1/Pik-2 and RGA5/RGA4 systems (36, 40, 41), our new results show the utility of structure-guided approaches to engineering NLR integrated domains to extend binding to different effectors. While recent advances in protein structure modelling will support future engineering efforts, challenges remain in the accurate prediction of side chain positions, and the effect of individual mutations, which will necessitate experimental determination of protein complexes to optimise intermolecular interactions.

The Pikp-1^SNK-EKE^ variant differs from the wild-type Pikh-1 allele in just two amino acid positions. Generating Pikp-1^SNK-EKE^ from Pikh-1 requires a maximum of four nucleotide substitutions, which can be achieved using precise base editing and prime editing technologies (49, 50). In many countries, edited crop varieties which do not contain DNA from another species are not subject to restrictions beyond those required for conventionally bred crop varieties. Therefore, this work raises the exciting prospect of editing wild-type alleles of NLRs that have greater potential for deployment in the field than those incorporating entirely new protein domains or substantial sequence changes.

Given the limited number of *M. oryzae* effectors for which host targets have been identified, it is notable that three (AVR-Pik, AVR- Pia and AVR1-CO39) interact with HMA domains of HIPPs and/or HPPs (8, 48). The AVR-Pik-like effector APikL2, which is highly conserved across *M. oryzae* lineages with different grass hosts, also binds to the HMA domain of a HIPP (sHMA94 from *Setaria italica*) (51). Although interaction with AVR-Pik appears to stabilise HIPPs (7), at present, the consequences and significance for pathogen virulence of these effector/HMA interactions are unclear. Intriguingly, the potato mop-top virus movement protein has been shown to interact with NbHIPP26 (52), and other HIPPs/HPPs have been described as host susceptibility factors (53–55) and may represent targets of effectors from other pathogens. This raises the possibility that incorporating different HMA domains into the Pik-1 chassis (with the mutations described here to prevent autoactivation, if necessary), could offer a suite of NLR proteins capable of recognising as-yet unknown effectors from diverse pathogens. Alongside biochemical approaches to identify effector-target interactions, advances in structural modelling could also enable identification of novel HMA-binding effectors. AVR-Pik, AVR-Pia and AVR1-CO39 all share the conserved MAX structural fold (56), and effectors with a similar structural core may bind HMAs. For example, an integrated HMA domain has been engineered to respond to the MAX effector AVR-Pib (41), although it is yet to be demonstrated that this effector binds host HMA targets. Identification of specific effector/HMA pairs could guide Pik-1 engineering for recognition of new pathogens, and potentially enable the design of synthetic HMA domains capable of binding to and recognising a broad range of pathogen effectors.

Advances in our understanding of the molecular and structural basis of NLR activation have progressed efforts for rational engineering of NLR proteins with altered recognition profiles. Previous studies have successfully engineered integrated domains to extend recognition capacities, though so far this has either resulted in regeneration of resistance already conferred by other NLRs (36, 40, 41) or provided recognition of a protein not present in the native pathogen (28). In this study, we use an effector target to guide engineering of an integrated domain to deliver two engineered Pik-1 variants with new-to-nature effector recognition profiles. The chimeric NLR Pikp-1^OsHIPP19-mbl7^ highlights the potential to incorporate diverse HMA domains without rendering the chimera autoactive. The triple mutant Pikp-1^SNK- EKE^ illustrates the benefit of structural/biochemical characterisation of effector-target interactions to inform rational engineering. Crucially, both engineered NLR proteins deliver novel resistance in transgenic rice, and have potential for deployment in the field against *M. oryzae* isolates carrying the stealthy AVR-PikC and AVR-PikF alleles. This study demonstrates the value of target-guided approaches in engineering NLR proteins with new-to-nature recognition profiles. We propose that this approach could expand the “toolbox” of resistance genes to counter the devastating impacts of plant pathogens on crop yields and global food security.

## Acknowledgments

This work was supported by the UKRI Biotechnology and Biological Sciences Research Council (BBSRC) Norwich Research Park Biosciences Doctoral Training Partnership, UK [grant BB/M011216/1]; the UKRI BBSRC, UK [grants BB/P012574, BBS/E/J/000PR9795, BB/M02198X], the European Research Council [ERC; proposal 743165]; The Thailand Research Fund through The Royal Golden Jubilee Ph.D. Program [PHD/0152/2556]; the John Innes Foundation; the Gatsby Charitable Foundation; The British Society for Plant Pathology (undergraduate vacation bursary); JSPS KAKENHI 15H05779 and 20H05681; JSPS/The Royal Society Bilateral Research for the project “Retooling rice immunity for resistance against rice blast disease” (2018–2019). We would also like to thank Julia Mundy and David Lawson from the JIC Biophysical Analysis and X-ray Crystallography platform for their support with protein crystallization and X-ray data collection, Andrew Davies and Phil Robinson from JIC Scientific Photography for their help with leaf imaging, Gerhard Saalbach and Carlo de Oliveira Martins from the JIC Proteomics platform for intact mass spectrometry analysis, and Dan Maclean from The Sainsbury Laboratory for support with statistical analyses. We also thank all members of the Banfield, Kamoun, and Terauchi groups for discussions.

## Materials and Methods

### Gene cloning for protein expression in *N. benthamiana*

For protein expression in planta, full length Pikp-1 NLRs containing the OsHIPP19 HMA domain (and Pikp-1^OsHIPP43-mbl7^), the Pikp-1^SNK-EKE^ mutation (made by introducing the S258E mutation in the HMA domain by PCR), and other HMA domain mutations were assembled using Golden Gate cloning into the plasmid pICH47742 with a C-terminal 6xHis/3xFLAG tag. Expression was driven by the *A. tumefaciens* Mas promoter and terminator. Full-length wild-type Pikp-1, Pikp-1^NK-KE^, Pikp-2, and AVR-Pik variants used were generated as described previously (9, 32, 33, 35, 36). All DNA constructs were verified by sequencing.

### Gene cloning, expression, and purification of proteins for in vitro binding studies

For SPR, Pikp-HMA^NK-KE^ and Pikp-HMA^SNK-EKE^ (residues Gly186 – Asp264) variants were cloned into pOPIN-M (generating a 3C protease cleavable N-terminal 6xHis:MBP-tag). AVR-PikD, AVR-PikC and AVR-PikF (residues Glu22 – Phe93) were cloned into pOPIN-E (generating a C-terminal non-cleavable 6xHis-tag, but also including a 3C protease cleavable N-terminal SUMO-tag, as detailed previously (9). The Pikp-HMA^NK-KE^ and Pikp-HMA^SNK-EKE^ proteins were expressed and purified using the same pipeline as described below for obtaining protein complexes for crystallisation, whereas the effectors were retained on the second pass through the 5 ml Ni^2+^-NTA column (which served to remove the SUMO tag following 3C cleavage) requiring specific elution with elution buffer (50 mM Tris-HCl pH 8.0, 50 mM glycine, 0.5 M NaCl, 500 mM imidazole, 5% (v/v) glycerol), followed by gel filtration using a Superdex 75 26/600 column equilibrated in running buffer (20 mM HEPES pH 7.5 and 150 mM NaCl). Proteins were concentrated and stored at -80 °C for further studies.

### Cloning, expression, and purification of proteins for crystallization

For crystallization of the Pikp-HMA^NK/KE^/AVR-PikC, Pikp-HMA^SNK/EKE^/AVR-PikC and Pikp- HMA^SNK/EKE^/AVR-PikF complexes, Pikp-HMA (residues Gly186 – Asp264) variants were cloned into pOPIN-M and AVR-PikC or AVR-PikF into pOPIN-A using InFusion cloning. Chemically competent *E. coli* SHuffle cells (57) were co-transformed with these vectors to produce 6xHis-MBP-tagged Pikp-HMA domains and untagged effectors. Cultures of these cells were grown in auto-induction media (58) to an OD600 of 0.4 – 0.6 at 30 °C, then incubated overnight at 18 °C. Cells were harvested by centrifugation and resuspended in lysis buffer (50 mM Tris-HCl pH 8.0, 50 mM glycine, 0.5 M NaCl, 20 mM imidazole, 5% (v/v) glycerol, 1 cOmplete^TM^ EDTA-free protease inhibitor cocktail tablet (Roche) per 50 ml buffer). Resuspended cells were lysed by sonication with a VibraCell sonicator (SONICS), and whole-cell lysate was clarified by centrifugation. An AKTA Xpress (GE Healthcare) system was used to carry out a two-step purification at 4 °C. The clarified cell lysate was first injected onto a 5 ml Ni^2+^-NTA column (GE Healthcare). The complexes were step-eluted with elution buffer (50 mM Tris-HCl pH 8.0, 50 mM glycine, 0.5 M NaCl, 500 mM imidazole, 5% (v/v) glycerol), and directly applied to a Superdex 75 26/600 gel filtration column equilibrated in running buffer (20 mM HEPES pH 7.5 and 150 mM NaCl). The 6xHis-MBP tag was cleaved from Pik-HMA by incubation with 3C protease (1 μg protease per mg of fusion protein) at 4 °C overnight and removed by passing the sample through a 5 ml Ni^2+^-NTA column connected to a 5 ml dextrin sepharose (MBPTrap) column (GE Healthcare), both equilibrated in lysis buffer. Fractions containing the relevant complexes were then concentrated and injected onto a Superdex 75 26/600 column equilibrated in running buffer. Eluted fractions containing protein complexes were concentrated to 13 mg/mL (Pikp-HMA^NK/KE^/AVR-PikC), 10 or 20 mg/mL (Pikp- HMA^SNK/EKE^/AVR-PikC) and 20 mg/mL (Pikp-HMA^SNK/EKE^/AVR-PikF) for crystallisation. Protein concentrations were determined using a Direct Detect Infrared Spectrometer (Millipore Sigma).

### Protein crystallisation, data collection, structure solution, refinement and validation

Crystallisation trials were set up in 96-well plates using an Oryx Nano robot (Douglas Instruments) with 0.3 μl of protein combined with 0.3 μl reservoir solution. Crystals of each complex were obtained in multiple conditions using the commercially available Morpheus® screen (Molecular Dimensions). Crystals used for Xray data collection were obtained from condition F8 (Pikp-HMA^NK/KE^/AVR-PikC complex), D7 (Pikp-HMA^SNK/EKE^/AVR-PikC), and H4 (Pikp-HMA^SNK/EKE^/AVR-PikF). The crystals were snap frozen in liquid nitrogen and shipped to Diamond Light Source for X-ray data collection. Diffraction data were collected at the Diamond Light Source, i04 and i03 beamlines (see table S1), under proposals mx13467 and mx18565. The data were scaled and merged by Aimless in the CCP4i2 software package (59). Each of the structures were solved by molecular replacement using PHASER (60). The search models used were the Pikp-HMA/AVR-PikD complex (PDB entry: 5A6W) for Pikp-HMA^NK/KE^/AVR- PikC, the Pikp-HMA^NK/KE^/AVR-PikC complex (PDB entry: 7A8W) for Pikp-HMA^SNK/EKE^/AVR- PikC, and the Pikp-HMA/AVR-PikD complex (PDB entry: 5A6W) for Pikp-HMA^SNK/EKE^/AVR- PikF. Iterative cycles of manual model building using COOT (61) and refinement with REFMAC (62) were used to derive the final structures, which were validated using the tools in COOT and MolProbity (63). The final protein structures, and the data used to derive them, have been deposited at the Protein Data Bank with IDs 7A8W (Pikp-HMA^NK/KE^/AVR-PikC), 7QPX (Pikp-HMA^SNK/EKE^/AVR-PikC), and 7QZD Pikp-HMA^SNK/EKE^/AVR-PikF).

### In vitro protein–protein interaction studies: Surface Plasmon Resonance (SPR)

Surface plasmon resonance was performed using a Biacore T200 (Cytiva) at 25 °C and at a flow rate of 30 µl/minute. The running buffer was 20 mM HEPES pH 7.5, 860 mM NaCl and 0.1%(v/v) Tween®20. Flow cell (FC) 2 of an NTA chip (GE Healthcare) was activated with 30 µl 0.5 mM NiCl2. 30 µl of the 6xHis-tagged effector (the ligand) was immobilised on FC2 to give a response of ∼250 RU. The HMA domain (the analyte) was then flowed over both FC1 and FC2 for 360 s, followed by a dissociation time of 180 s. Three separate concentrations of each HMA were tested, 4 nM, 40nM and 100nM. The NTA chip was regenerated after each cycle with 30 µl 0.35 M EDTA pH 8.0. The background response from FC1 (non-specific binding of the HMA domain to the chip) was subtracted from the response from FC2. To obtain %Rmax, the binding response (Robs) was measured immediately prior to the end of injection and expressed as a percentage of the theoretical maximum response (Rmax) assuming a 2:1 HMA:effector binding model for Pikp-HMA, Pikp^NK-KE^-HMA and Pikp^SNK-EKE^-HMA calculated as follows:

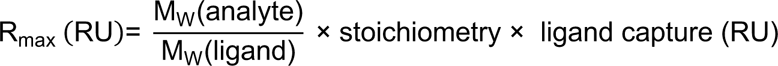

Data analysis and visualisation was carried out in R v4.1.2 (64) using the packages dplyr (v1.0.9 (65)) and ggplot2 (v3.3.6 (66)).

### *N. benthamiana* cell death assays

*A. tumefaciens* GV3101 (C58 (rifR) Ti pMP90 (pTiC58DT-DNA) (gentR) Nopaline(pSouptetR)) cells carrying relevant Pikp-1 constructs, were resuspended in agroinfiltration media (10 mM MES pH 5.6, 10 mM MgCl2 and 150 μM acetosyringone) and mixed with *A. tumefaciens* GV3101 carrying Pikp-2, AVR-Pik effectors, and P19 at OD600 0.4, 0.4, 0.6 and 0.1, respectively. 4-week old *N. benthamiana* leaves were infiltrated using a needleless syringe.

*N. benthamiana* plants were grown in a controlled environment room at 22°C constant temperature and 80% relative humidity, with a 16 hour photoperiod, and were returned to the same room following infiltration. Leaves were collected at 5 dpi (days post infiltration) and photographed under visible and UV light. Images shown are representative of three independent experiments, with a minimum of ten repeats (leaves) in each experiment. The cell death index used for scoring is as presented previously (32). Data analysis and visualisation was carried out in R v4.1.2 (64) using the packages dplyr (v1.0.9 (65)) and ggplot2 (v3.3.6 (66)). Relevant comparisons between conditions (i.e. combinations of NLRs and effectors) in cell death assays were made using estimation methods (67) using the package besthr (v0.2.0 (68)). Mean scores were calculated for each condition and each biological replicate. These means were ranked, and a mean rank calculated for each condition. Bootstrapping was then used to estimate the confidence interval of the mean rank of each condition. 1000 samples were drawn (with replacement) from the values present in the dataset, and the mean rank calculated for each of these samples to give a distribution of mean rank estimates. The 2.5 and 97.5 quantiles were determined. Conditions are considered to be different if their means lie outside the 2.5 and 97.5 quantiles.

### Confirmation of protein production in cell death assays by Western blot analysis

Western blot analysis was used to confirm the presence of proteins in *N. benthamiana* during cell death assays. 3 leaf discs were taken 2dpi, flash frozen in liquid nitrogen, and ground to a fine powder using a micropestle. Leaf powder was mixed with 300 μl of plant protein extraction buffer (GTEN (25 mM Tris-HCl, pH 7.5, 150 mM NaCl, 1 mM EDTA, 10 % v/v glycerol), 10 mM DTT, 2 % (w/v) PVPP, 0.1 % Tween®-20, 1x plant protease inhibitor cocktail (Sigma)). The sample was clarified by centrifugation (20,000 x *g* for 2 minutes at 4°C, twice) and 40μL of the resulting supernatant added to 10 μL SDS-PAGE loading dye (RunBlue^TM^ 4x LDS sample buffer (Expedeon)), followed by incubation at 95°C for 5 minutes.

Samples were subjected to SDS-PAGE/western blot analysis to detect epitope-tagged proteins. Pikp-1, Pikp-2 and AVR-Pik effectors were detected by probing membranes with anti-FLAG-HRP (Generon, 1:10000 dilution), anti-HA-HRP (ThermoFisher Scientific, 1:3000 dilution) and anti-Myc-HRP (Santa Cruz, 1:5000 dilution) antibodies, respectively, and LumiBlue ECL Extreme (Expedeon). Membranes were also stained with Ponceau S to observe protein loading.

### In planta protein-protein interaction studies: co-immunoprecipitation

For co-immunoprecipitation assays, 3 leaves were harvested 3 dpi and flash-frozen in liquid nitrogen. Leaf tissue was ground to a fine powder in liquid nitrogen using a pre-chilled pestle and mortar and resuspended in ice-cold plant protein extraction buffer (2ml / mg of powder). Plant cell debris was pelleted by centrifugation at 4,200 x *g* for 30 minutes at 4 °C and the supernatant was filtered through a 0.45 μm membrane. 20 μL of filtered extracts were combined with 5 μL SDS-PAGE loading dye as input samples. For immunoprecipitation, 1 mL of filtered protein extract was mixed with 40μL of anti-FLAG M2 magnetic beads (Merck, formerly Sigma-Aldrich) (equilibrated in GTEN + 0.1 % Tween-20 prior to use) in a rotary mixer for 1 hr at 4 °C. The beads were washed 5X in ice-cold IP buffer by separating the beads using a magnetic rack, and the proteins eluted by resuspending the beads in 30 μL SDS loading buffer and incubating at 70 °C for 10 minutes. Following SDS-PAGE and transfer, membranes were probed with antibodies as above.

### Fungal strains and transformation

To generate *M. oryzae* Sasa2 harboring different *AVR-Pik* alleles or *AVR-Pii*, Sasa2 was transformed individually with expression vectors for AVR-PikC, -PikD and -PikF and AVR-Pii. The expression vectors used for generating transgenic *M. oryzae* Sasa2 were pCB1531:*AVR-Pii* promoter:*AVR-Pii* constructed by Yoshida et al. (34), pCB1531:*AVR-PikD* promoter:*AVR-PikC*, -PikD constructed by Kanzaki et al. (30), and pCB1531:*AVR-PikD* promoter:*AVR-PikF*, generated according to (30). The template DNA for AVR-PikF was synthesized by GENEWIZ (Genewiz, Saitama, Japan). These expression vectors were used to transform Sasa2 (lacking *AVR-PikD* alleles and *AVR-Pii*) following the method of Sweigard et al. (69).

### Rice transformation and confirmation of transgene expression

To generate constructs for rice transformation, Golden Gate Level 1 constructs encoding a hygromycin resistance cassette (35S promoter/nos terminator), untagged *Pikp-1/Pikp- 1^OsHIPP19-mbl7^/Pikp-1^SNK-EKE^* with the NLR flanked by the 35S promoter and terminator, and untagged *Pikp-2* also flanked by the 35S promoter and terminator were assembled by Golden Gate assembly into the Level 2 vector pICSL4723. The resulting Level 2 constructs contained the hygromycin resistance cassette, 35S::*Pikp-1/Pikp-1^OsHIPP19-mbl7^/Pikp-1^SNK-EKE^* and 35S::*Pikp-2*. These constructs were introduced into *Agrobacterium tumefaciens* (strain EHA105) and used for Agrobacterium-mediated transformation of *Oryza sativa* cv. Nipponbare following the method of Okuyama et al. (37).

PCR confirmation of the presence of *Pikp-1/Pikp-2* transgenes in T1 progenies used two primer sets, *Pikp-1* (F:TGATCAAAGACCACTTCCGCGTTC + R:TGCTGCCCGCAATGTTTTCACTGC) and *Pikp-2* (F:ATTGTATATGTCAGCCAGAAAATG + R:TCCTCAGGGACTTGCTCGTCTAC) that amplify *Pikp-1* and *Pikp-2*, respectively.

To confirm transgene expression, total RNA was extracted from leaves using an SV Total RNA Isolation System (Promega, WI, USA) and used for RT-PCR. cDNA was synthesized from 500 ng total RNA using a Prime Script RT Reagent Kit (Takara Bio, Otsu, Japan). RT-PCR was performed using three primer sets, *Pikp-1* (F:TGATCAAAGACCACTTCCGCGTTC

+ R:TGCTGCCCGCAATGTTTTCACTGC), *Pikp-2* (F:ATTGTATATGTCAGCCAGAAAATG +

R:TCCTCAGGGACTTGCTCGTCTAC) and *OsActin* (F: CTGAAGAGCATCCTGTATTG + R:

GAACCTTTCTGCTCCGATGG) that amplify *Pikp-1*, *Pikp-2* and *OsActin*, respectively.

### Disease resistance/virulence assays in rice

Rice leaf blade punch inoculation was performed using the *M. oryzae* isolates. A conidial suspension (3 × 10^5^ conidia mL^−1^) was punch inoculated onto a rice leaf one month after seed sowing. The inoculated plants were placed in a dark dew chamber at 27 °C for 24 h and then transferred to a growth chamber with a 16 h light/8 h dark photoperiod. Disease lesions were scanned at 7 days post-inoculation (dpi), and lesion size was measured manually using Image J software (70).

---

**Supplementary figure 1.**
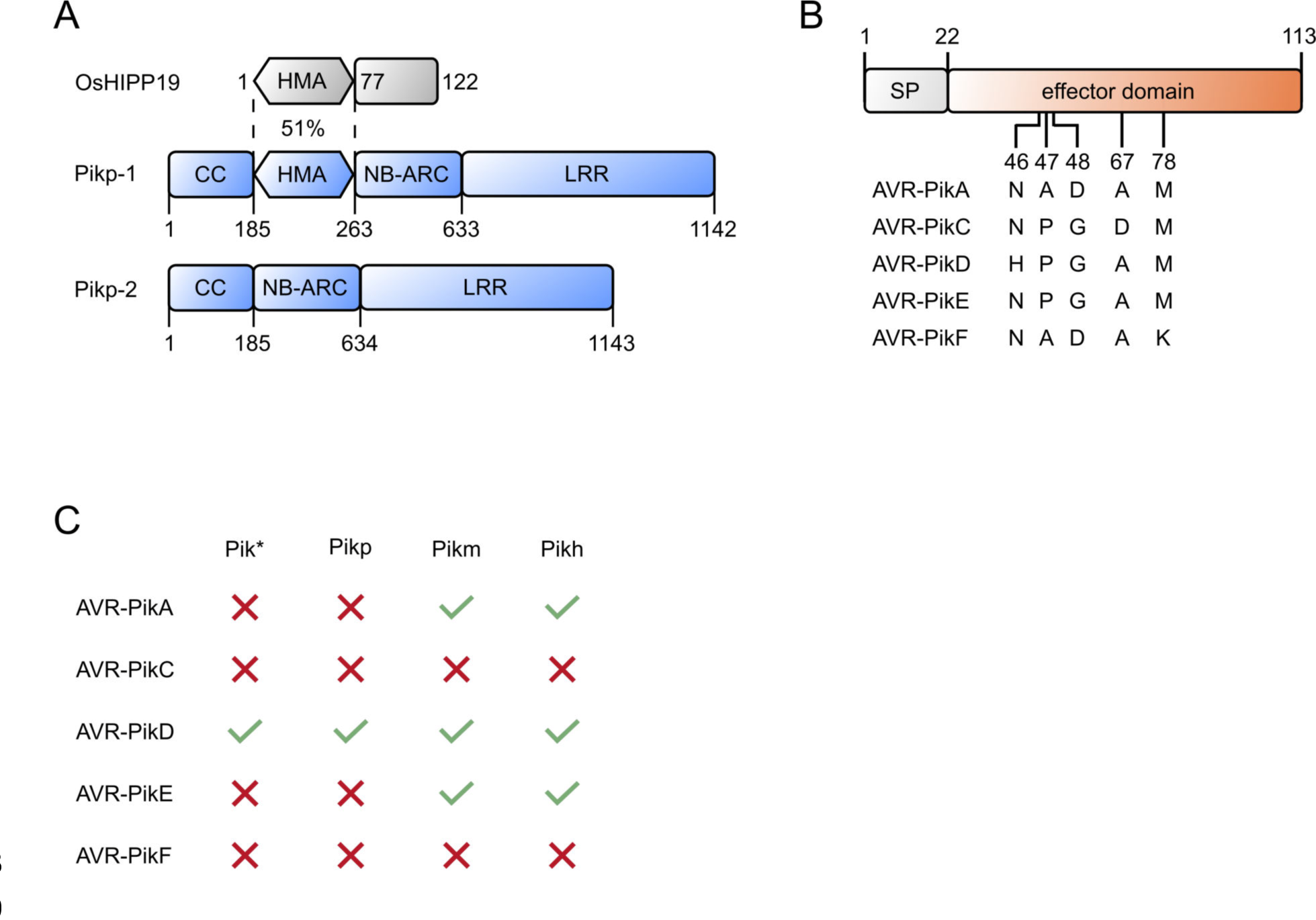
Domain structure and resistance profiles for the proteins in this study. (**A**) Cartoon representation of the OsHIPP19, Pik-1, and Pik-2 domains, with numbers giving the amino acid domain boundaries. CC = Coiled-coil, HMA = Heavy-metal-associated, NB-ARC = Nucleotide-binding found in APAF-1, R proteins and CED4, LRR = Leucine-rich repeat. (**B**) Cartoon representation of AVR-Pik effector variants. Amino acid polymorphisms between variants are shown as single letter codes with the number giving the position. SP = signal peptide. (**C**) Summary of the recognition profiles of known Pik NLR alleles against AVR-Pik variants (tick = resistant, cross = susceptible).

**Supplementary figure 2.**
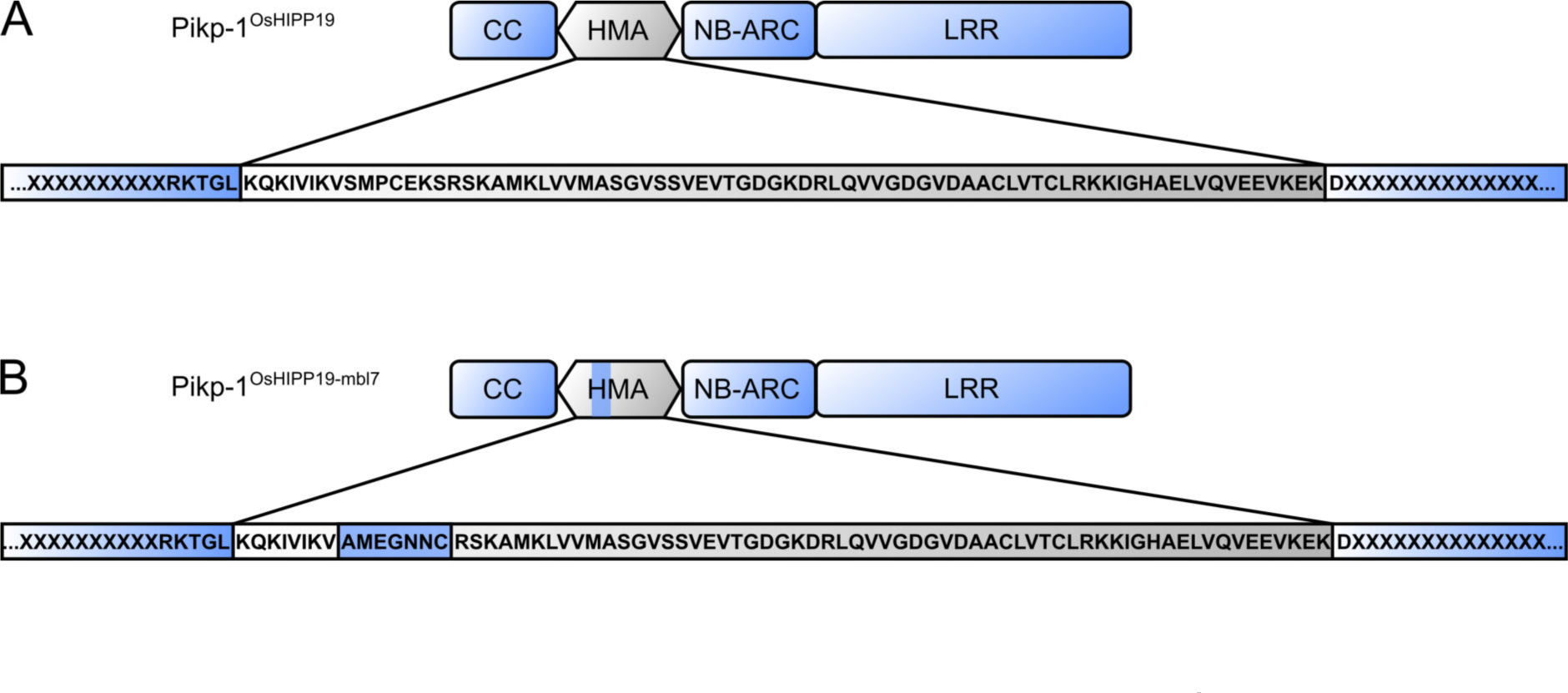
Schematic representation of the Pikp-1^OsHIPP19^ (A) and Pikp- 1^OsHIPP19-mbl7^ (B) chimeras. The amino acid sequence below indicates junctions between the sequence derived from Pikp-1 (blue highlight) and from OsHIPP19 (grey highlight).

**Supplementary figure 3.**
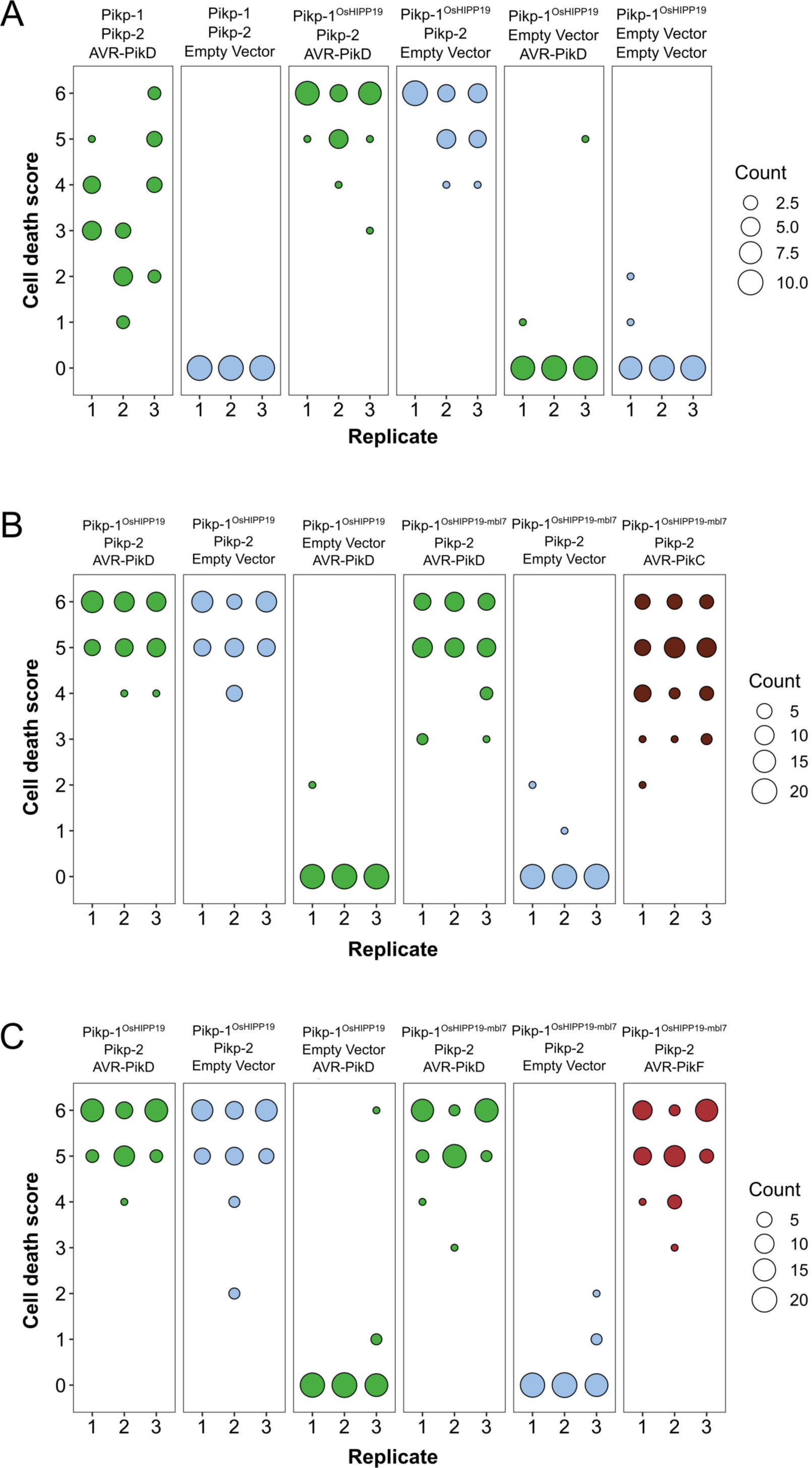
Pikp-mediated response scoring represented as dot plots, subdivided by replicate, for repeats of experiments presented in. Figure 1b**, 1d and 1f (panels A, B, and C; respectively).** Each replicate consisted of 10 (A) or 20 (B, C) repeats for each sample. Fluorescence intensity is scored as described in Figure 1. Scores from the three replicates in panels A, B, and C were combined and represented as the dot plots in Figure 1c, 1e and 1g, respectively.

**Supplementary figure 4.**
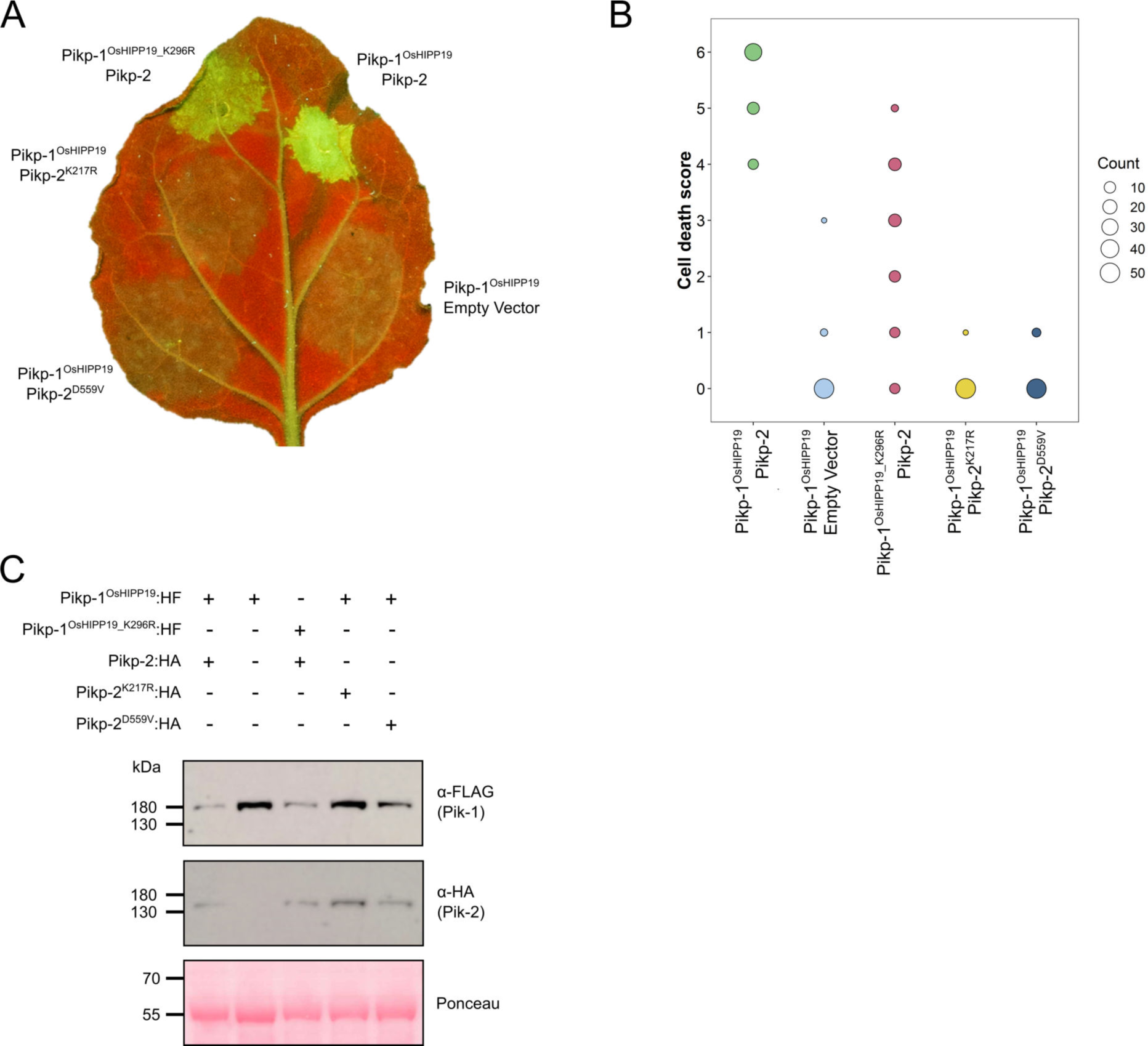
The Pikp-1^OsHIPP19^ chimera requires both the P-loop and MHD motifs in Pikp-2 for autoactivity, and the P-loop in Pikp-1 for full cell death. (**A**) Example leaf showing the P-loop mutant (K217R, middle, left) and MHD mutant (D559V, bottom, left) in Pikp-2 abolishes autoactive cell death, whereas the P-loop mutant in Pikp-1^OsHIPP19^ (K196R, top, left) reduces the cell death response. (**B**) Pikp-mediated response scoring represented as dot plots to summarise 54 repeats of the experiment shown in (**A**) across three independent experiments (Materials and Methods, figure S5). Fluorescence intensity is scored as stated in Figure 1. (**C**) Western blots confirming the accumulation of proteins in *N. benthamiana*. Plant cell lysates were probed for the expression of Pikp-1, Pikp-2, and mutants thereof, using anti-FLAG and anti-HA antiserum, respectively. Total protein extracts were visualised by Ponceau Staining.

**Supplementary figure 5.**
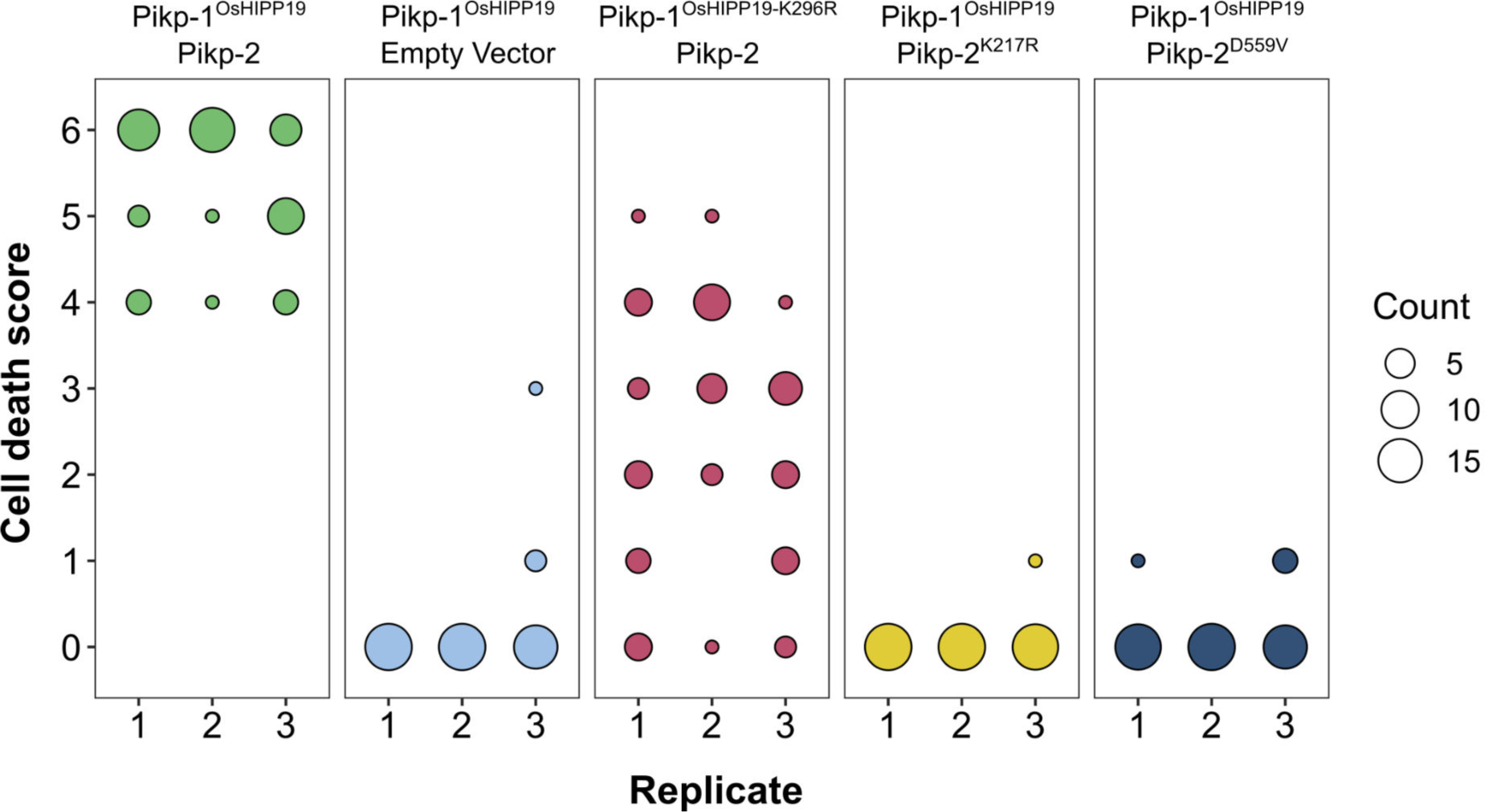
Pikp-mediated response scoring represented as dot plots, subdivided by replicate, for repeats of the experiment presented in Supplementary figure S3A. Each replicate consisted of 18 repeats for each sample. Fluorescence intensity is scored as described in Figure 1. Scores from the three replicates were combined and represented as the dot plot in Supplementary figure S4b.

**Supplementary figure 6.**
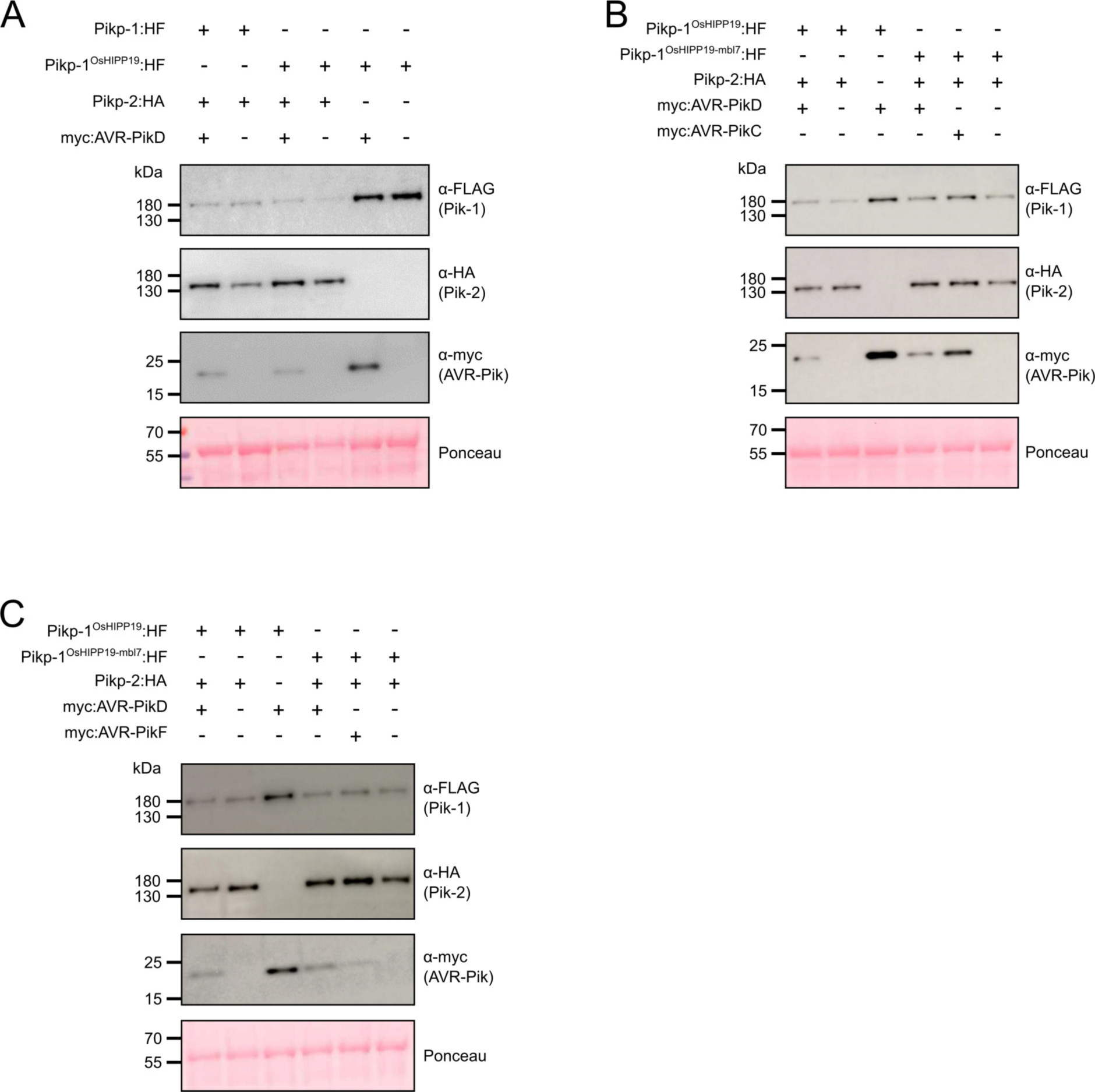
Western blots confirming the accumulation of proteins in *N. benthamiana* for the cell death assays shown in. Figure 1. Plant cell lysates were probed for the expression of Pikp-1/Pikp-1^OsHIPP19^/ Pikp-1^OsHIPP19-mbl7^, Pikp-2, and AVR-Pik effector variants using anti-FLAG, anti-HA and anti-Myc antiserum, respectively. Total protein extracts were visualised by Ponceau Staining.

**Supplementary figure 7.**
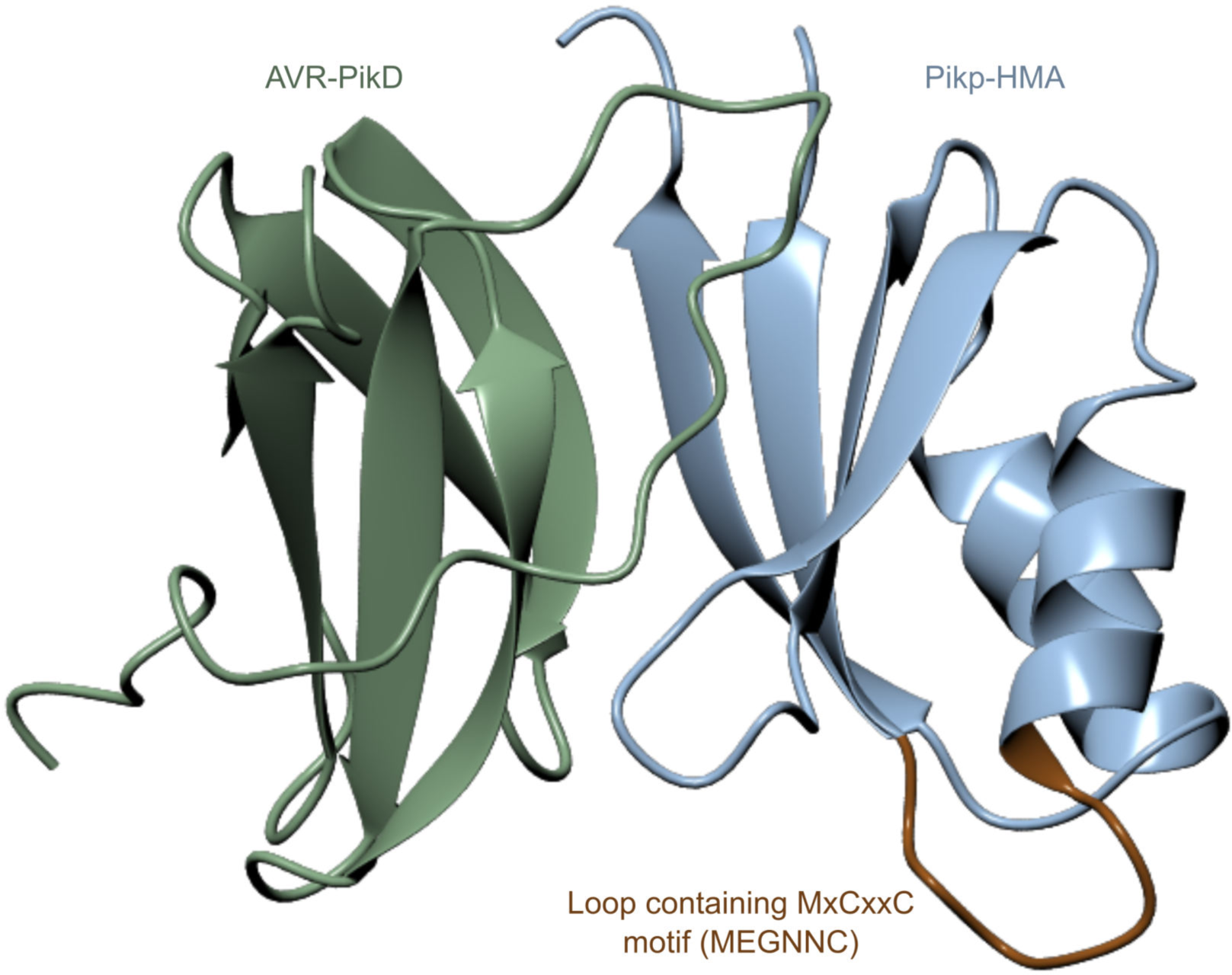
Location of the β1-α1 loop (brown) in Pikp-HMA (blue) is distant from the effector (green) binding surface in the crystal structure of complexes between these proteins. Structure shown is based on PDB entry 6G10. Protein structures are presented as ribbons.

**Supplementary figure 8.**
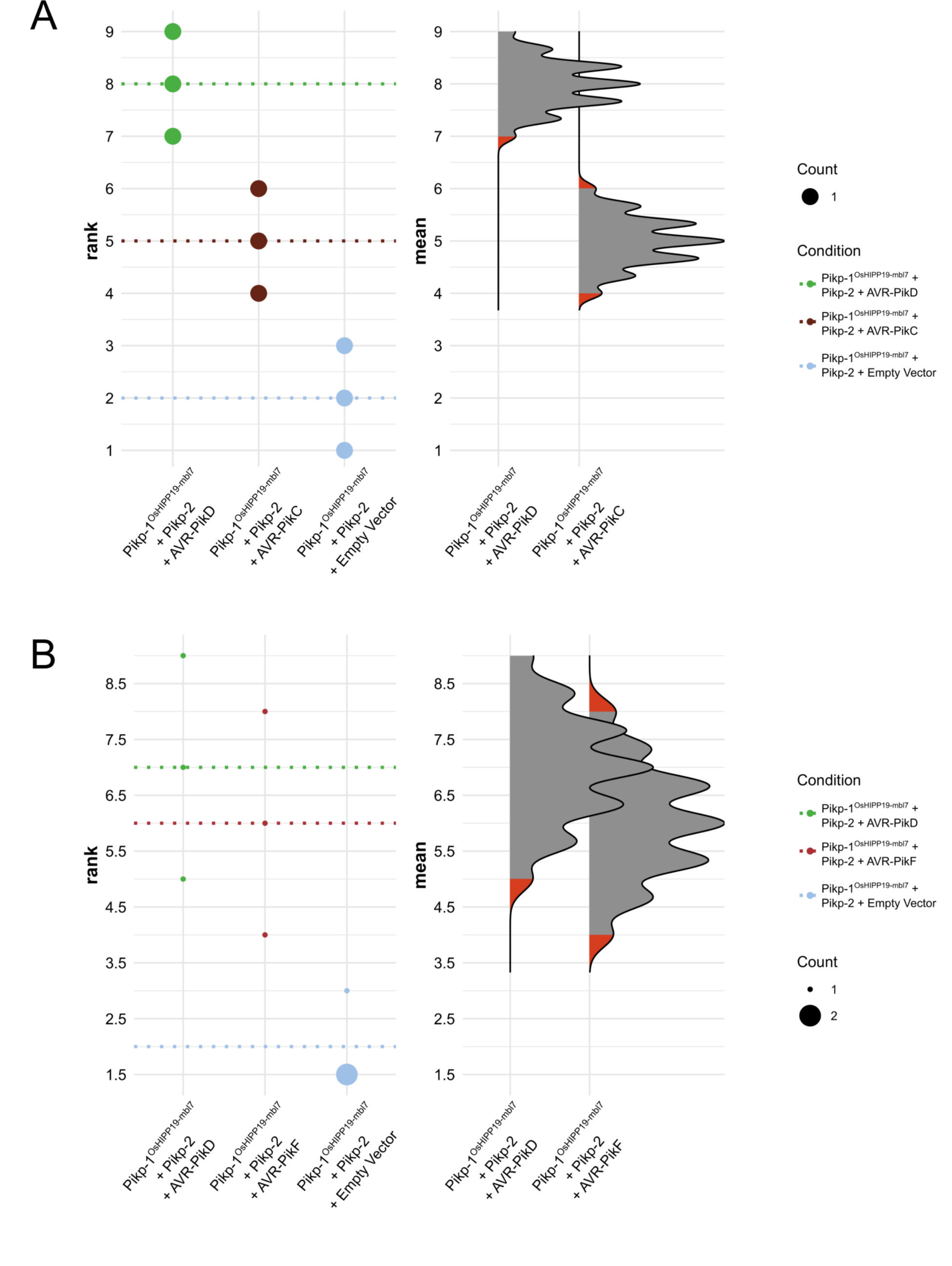
Statistical analysis by estimation methods of the cell death assays presented in Figure 1, for Pikp-1^OsHIPP19^/Pikp-2 with (A) AVR-PikD, AVR-PikC and empty vector, and (B) AVR-PikD, AVR-PikF and empty vector. **The panel on the left** represents the ranked data (dots) for the three replicates of each effector/control, and their corresponding mean (dotted line). The size of the dots is proportional to the number of observations with that specific value. The panel on the right shows the distribution of 1000 bootstrap sample rank means for Pikp-1^OsHIPP19^/Pikp-2/AVR-PikD and Pikp-1^OsHIPP19^/Pikp- 2/AVR-PikC (A) or Pikp-1^OsHIPP19^/Pikp-2 (B). The red areas represent the 2.5^th^ and 97.5^th^ percentiles of the distribution. The response of Pikp-1^OsHIPP19-mbl7^/Pikp-2 to AVR-PikD, AVR- PikC and AVR-PikF is considered significantly different to the response of Pikp-1^OsHIPP19- mbl7^/Pikp-2 to the empty vector as the rank mean of the latter (dotted line, left panel) falls beyond the red regions of the Pikp-1^OsHIPP19-mbl7^/Pikp-2/AVR-PikD, Pikp-1^OsHIPP19-mbl7^/Pikp- 2/AVR-PikC, and Pikp-1^OsHIPP19-mbl7^/Pikp-2/AVR- mean distributions.

**Supplementary figure 9.**
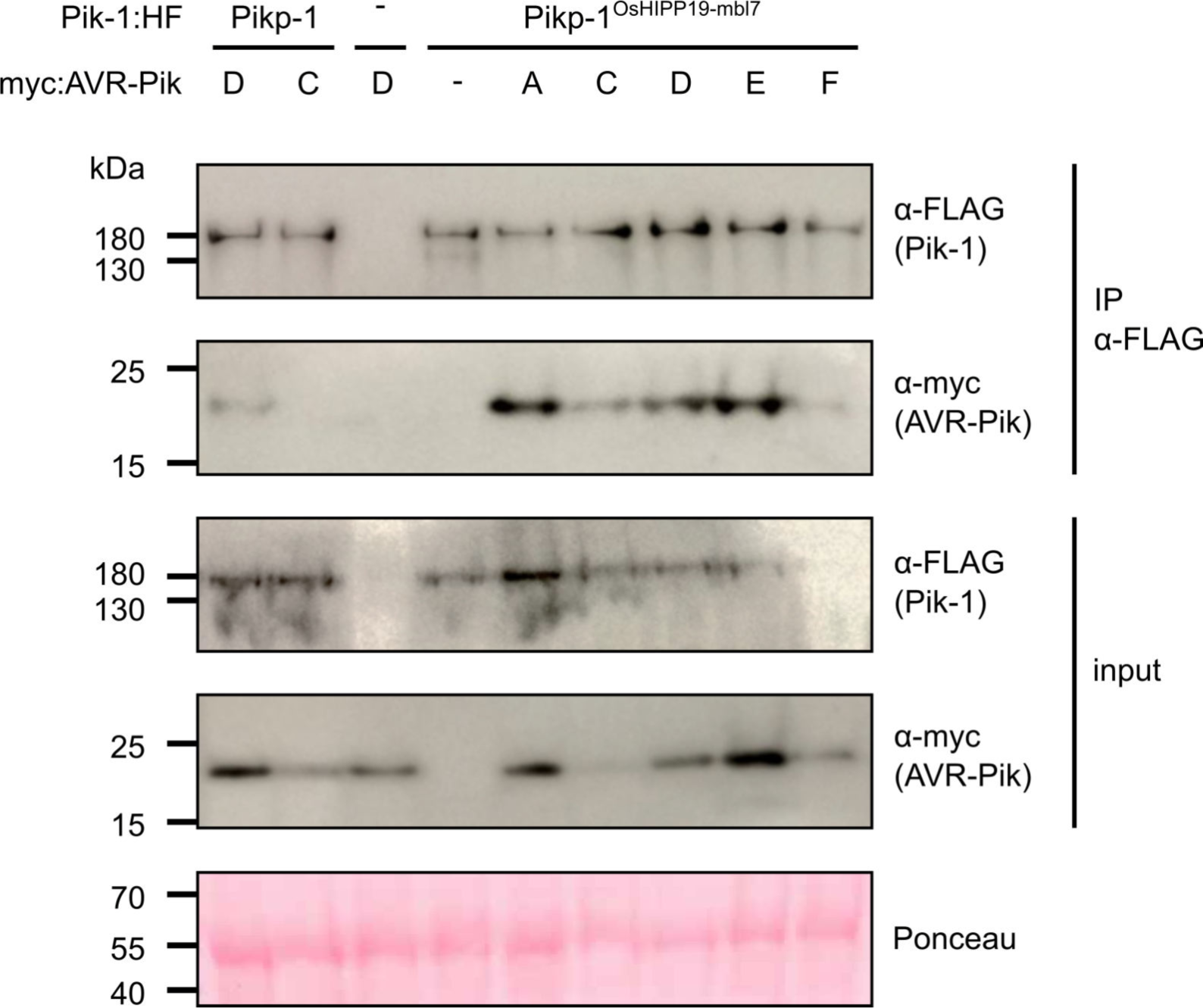
Western blots following co-immunoprecipitation show that the Pikp-1^OsHIPP19-mbl7^ chimera retains binding to all AVR-Pik effector variants in *N. benthamiana*. Plant cell lysates were probed for the expression of Pikp-1/Pikp-1^OsHIPP19-mbl7^ and AVR-Pik effector variants using anti-FLAG and anti-Myc antiserum, respectively. Total protein extracts were visualised by Ponceau Staining.

**Supplementary figure S10.**
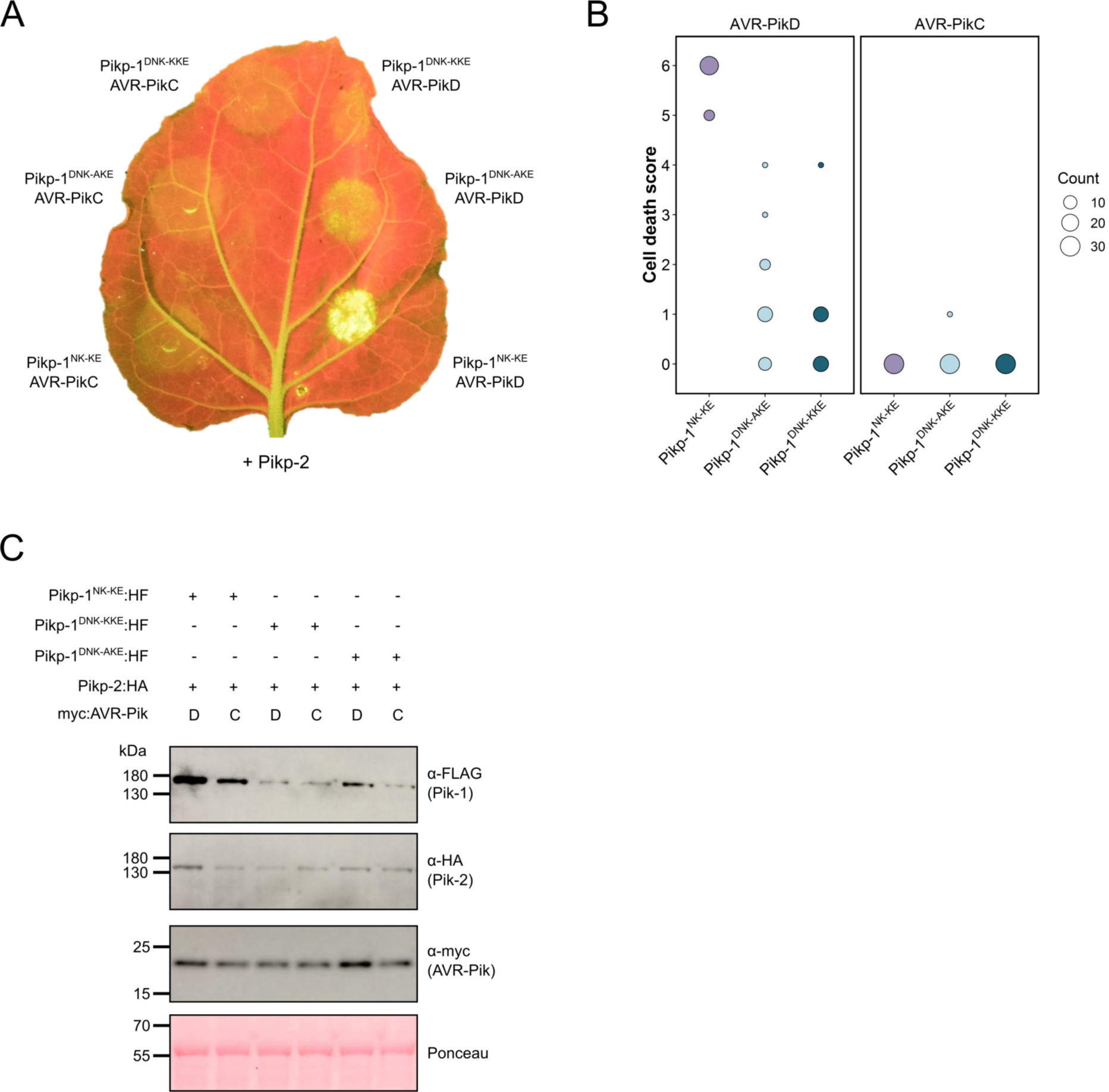
The mutations D224A and D224K mutations in the Pikp^NK-KE^ background do not extend response to AVR-PikC. (**A**) Neither the Pikp-1^DNK-AKE^ nor the Pikp-1^DNK-KKE^ mutant gains response to AVR-PikC (left, middle and left, top) and response to AVR-PikD is reduced in both mutants (right, middle and right, top). All infiltration spots contain Pikp-2. (**B**) Pikp-mediated response scoring represented as dot plots to summarise 30 repeats of the experiment shown in (**A**) across three independent experiments (Materials and Methods, figure S11). Fluorescence intensity is scored as stated in Figure 1. (**C**) Western blots confirming the accumulation of proteins in *N. benthamiana*. Plant cell lysates were probed for the expression of Pikp-1, Pikp-2, and effector variants, using anti-FLAG, anti-HA and anti-Myc antiserum, respectively. Total protein extracts were visualised by Ponceau Staining.

**Supplementary figure 11.**
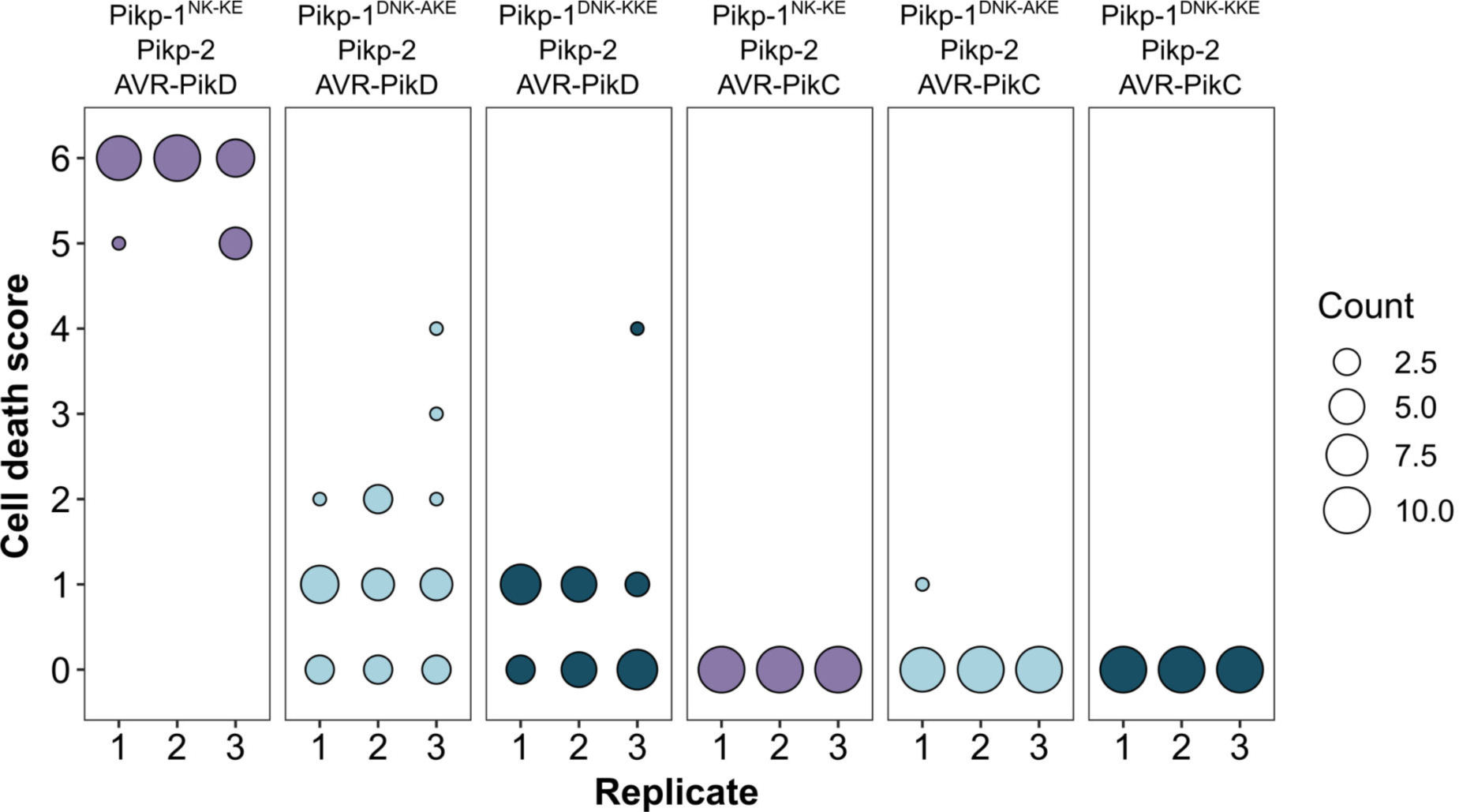
Pikp-mediated response scoring represented as dot plots, subdivided by replicate, for repeats of the experiment presented in Supplementary figure S7a. Each replicate consisted of 10 repeats for each sample. Fluorescence intensity is scored as described in Figure 1. Scores from the three replicates were combined and represented as the dot plot in Supplementary figure S10b.

**Supplementary figure 12.**
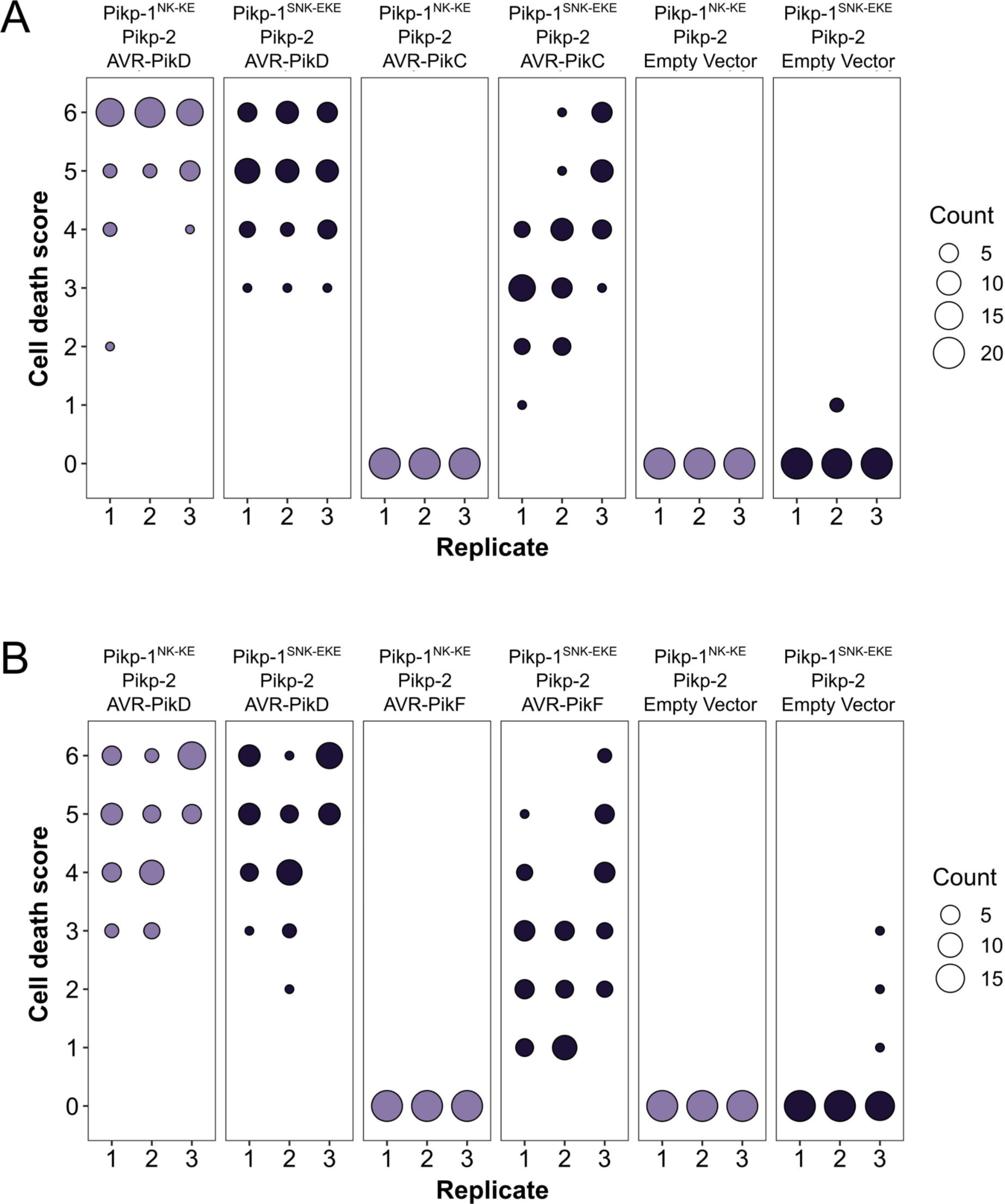
Pikp-mediated response scoring represented as dot plots, subdivided by replicate, for repeats of the experiment presented in Figures 2b (A) and 2d (B). Each replicate consisted of 20 (A) and 19 (B) repeats for each sample. Fluorescence intensity is scored as described in Figure 1. Scores from the three replicates were combined and represented as the dot plots in Figures 2c and 2e, respectively.

**Supplementary figure S13.**
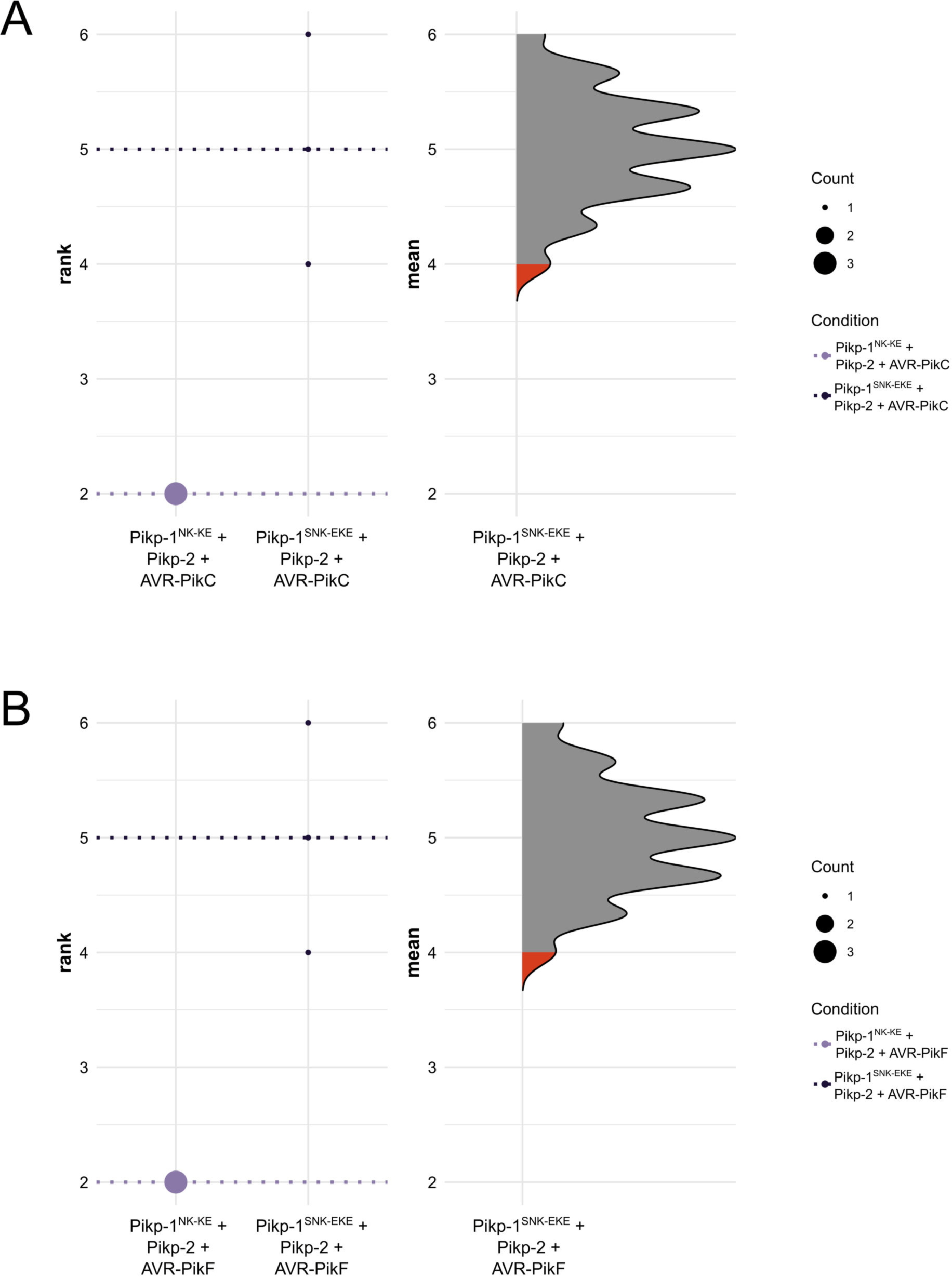
Statistical analysis by estimation methods of the cell death assays presented in Figure 2, for Pikp-1^NK-KE^/Pikp-2 and Pikp-1^SNK-EKE^/Pikp-2 with **(A)** AVR- AVR-PikC and **(B)** AVR-PikF. The panel on the left represents the ranked data (dots) for the three replicates of each receptor/effectorl, and their corresponding mean (dotted line). The size of the dots is proportional to the number of observations with that specific value. The panel on the right shows the distribution of 1000 bootstrap sample rank means for Pikp-1^SNK- EKE^/Pikp-2/AVR-PikC (A) or Pikp-1^SNK-EKE^/Pikp-2/AVR-PikF (B). The red areas represent the 2.5^th^ and 97.5^th^ percentiles of the distribution. The response of Pikp-1^SNK-EKE^/Pikp-2 to AVR- AVR-PikC/AVR-PikF is considered significantly different to the response of Pikp-1^NK-KE^/Pikp-2 to AVR-PikC/AVR-PikF as the rank mean of the latter (dotted line, left panel) falls beyond the red regions of the Pikp-1^NK-KE^/Pikp-2/AVR-PikC and Pikp-1^NK-KE^/Pikp-2/AVR-PikF mean distributions.

**Supplementary figure 14.**
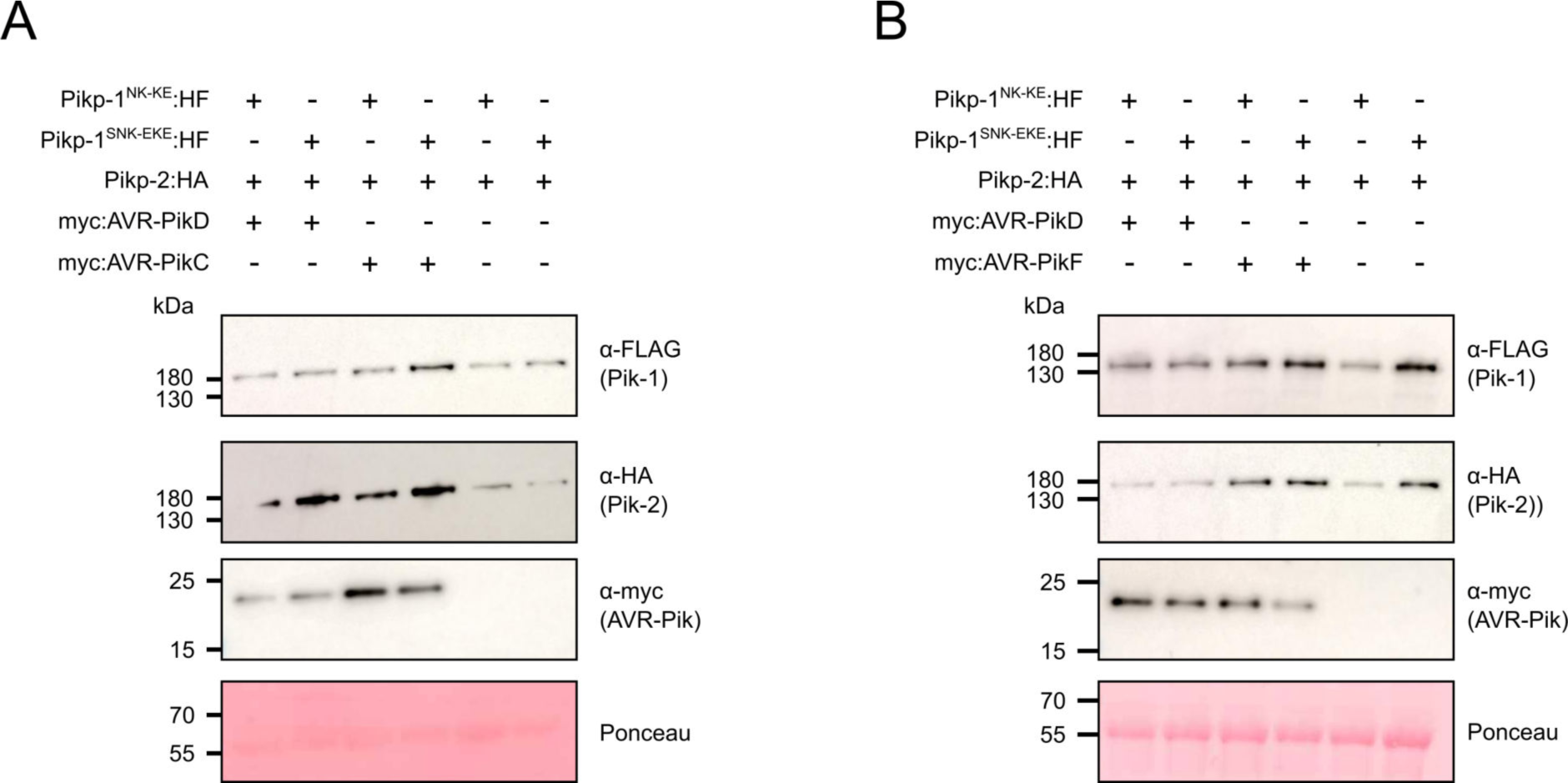
Western blots confirming the accumulation of proteins in *N. benthamiana* for the cell death assays shown in. Figure 2. (**A**) Accumulation of proteins for the experiments with AVR-PikC. (**B**) Accumulation of proteins for the experiments with AVR-PikF. Plant cell lysates were probed for the expression of Pikp-1^NK-KE^/Pikp-1^SNK-EKE^, Pikp-2, and AVR-Pik effector variants using anti-FLAG, anti-HA and anti-Myc antiserum, respectively. Total protein extracts were visualised by Ponceau Staining.

**Supplementary figure 15.**
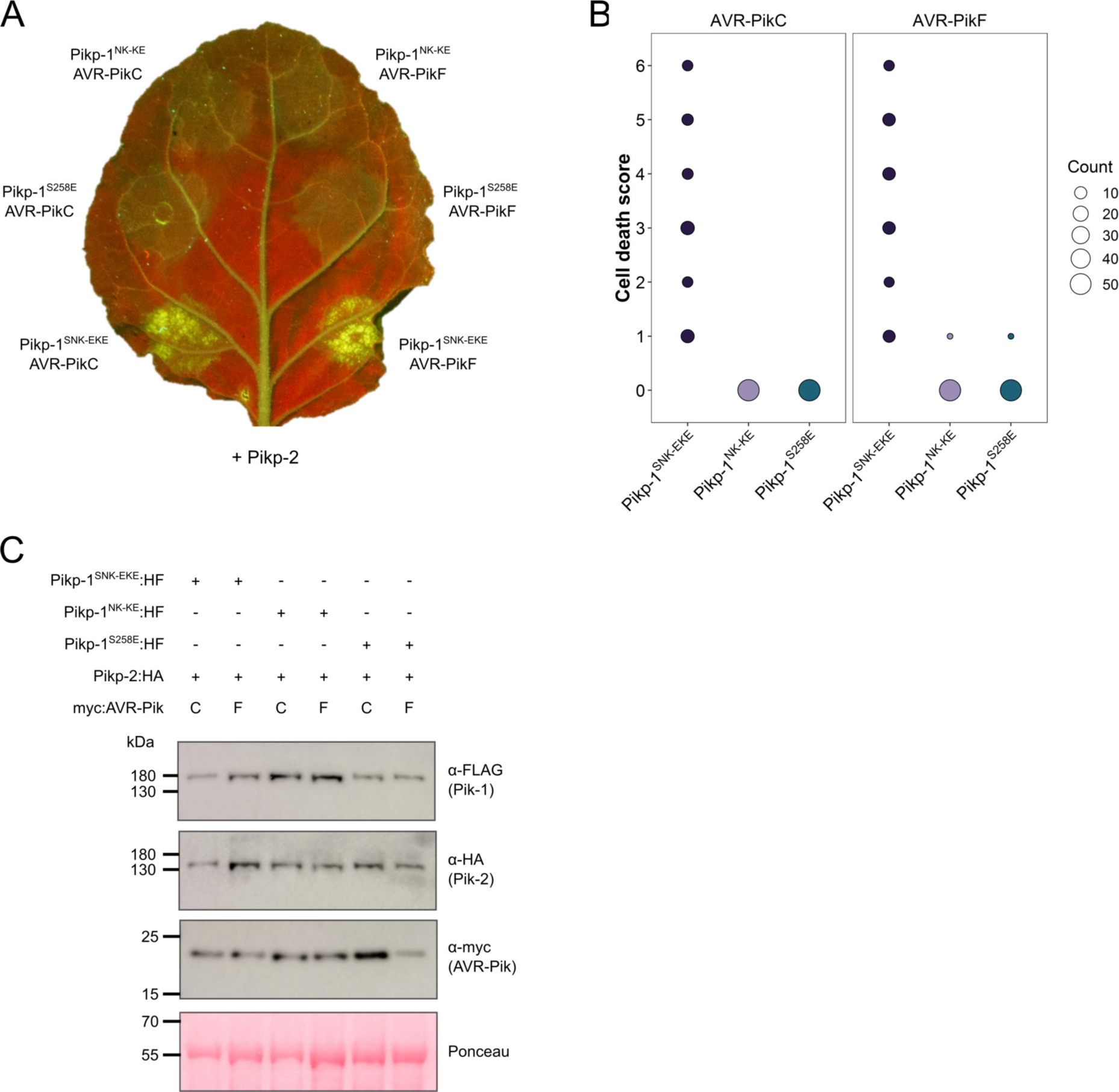
The Pikp S258E mutation alone does not extend response to AVR-PikC or AVR-PikF. (**A**) The Pikp-1^S258E^ mutant does not gain response to AVR-PikC (left, middle) or AVR-PikF (right, middle). All infiltration spots contain Pikp-2. (**B**) Pikp-mediated response scoring represented as dot plots to summarise 54 repeats of the experiment shown in (**A**) across three independent experiments (Materials and Methods, figure S16). Fluorescence intensity is scored as stated in Figure 1. (**C**) Western blots confirming the accumulation of proteins in *N. benthamiana*. Plant cell lysates were probed for the expression of Pikp-1, Pikp-2, and effector variants, using anti-FLAG, anti-HA and anti-Myc antiserum, respectively. Total protein extracts were visualised by Ponceau Staining.

**Supplementary figure 16.**
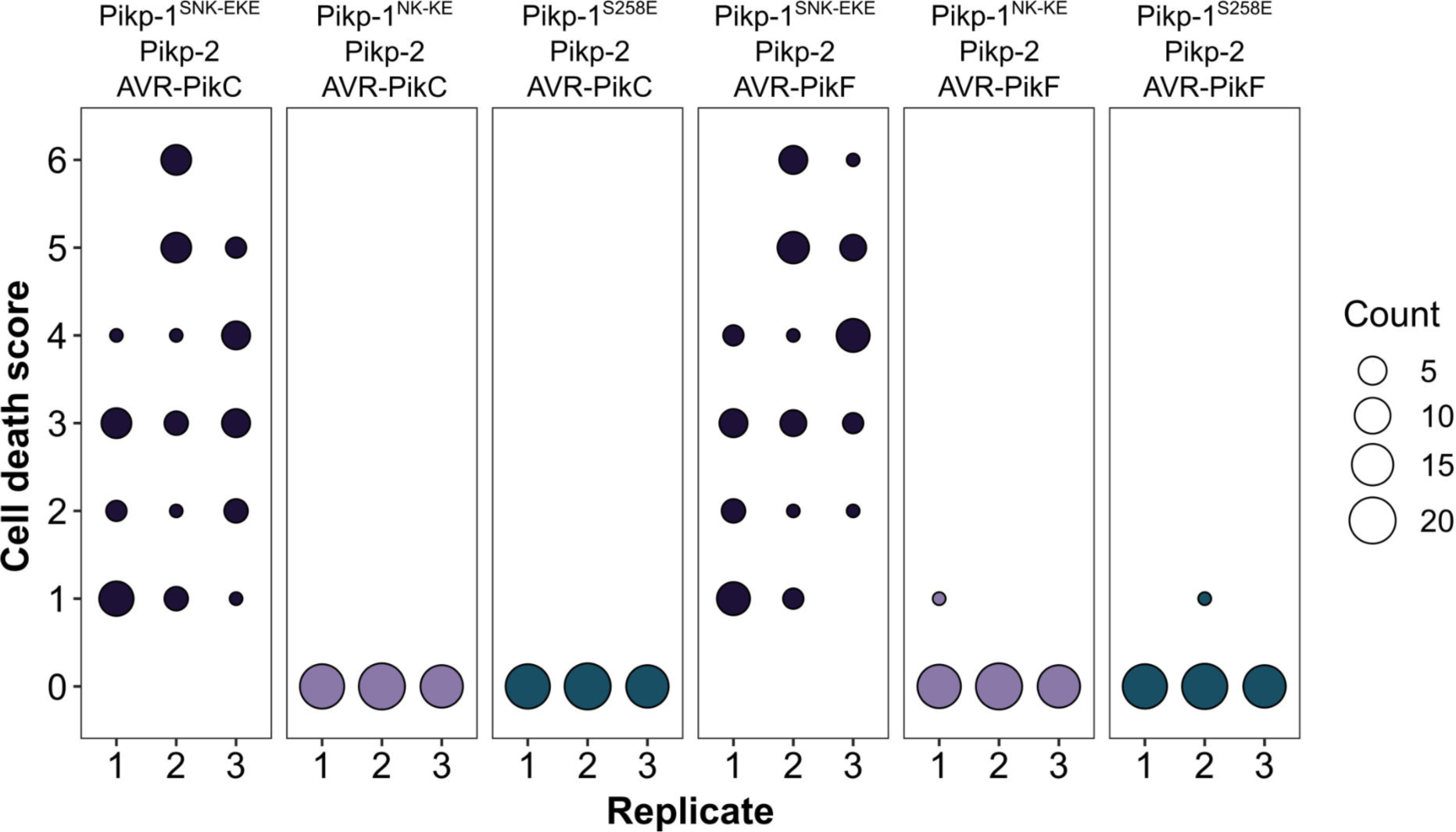
Pikp-mediated response scoring represented as dot plots, subdivided by replicate, for repeats of the experiment presented in Supplementary figure 15. The three replicates consisted of 18, 20 and 16 repeats for each sample, respectively. Fluorescence intensity is scored as described in Figure 1. Scores from the three replicates were combined and represented as the dot plot in Supplementary figure 15b, respectively.

**Supplementary figure 17.**
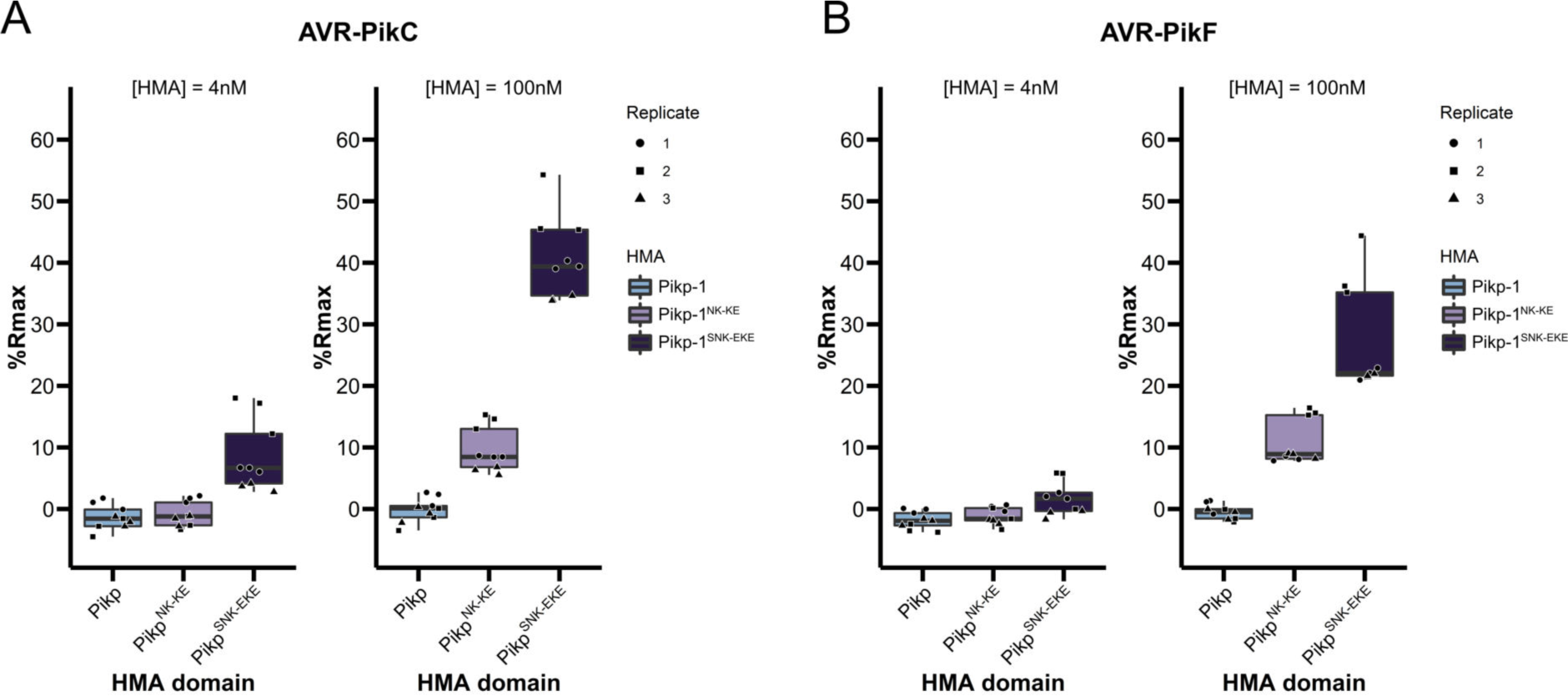
Boxplots showing the %Rmax observed for the interactions between AVR-PikC (**A**) or AVR-PikF (**B**), both at 4nM and 100nM injection concentrations, and each of Pikp-HMA, Pikp-HMA^NK-KE^ and Pikp-HMA^SNK-EKE^. %Rmax is the percentage of the theoretical maximum response, assuming a 2:1 binding model (as previously observed for Pikp-HMA proteins, see Materials and Methods).The center line of the box represents the median and the box limits are the upper and lower quartiles. The whiskers extend to the smallest value within Q1 − 1.5Å∼ the interquartile range (IQR) and the largest value within Q3 + 1.5Å∼ IQR. Individual data points are represented as black shapes. The experiment was repeated three times, with each experiment consisting of three technical replicates.

**Supplementary figure 18.**
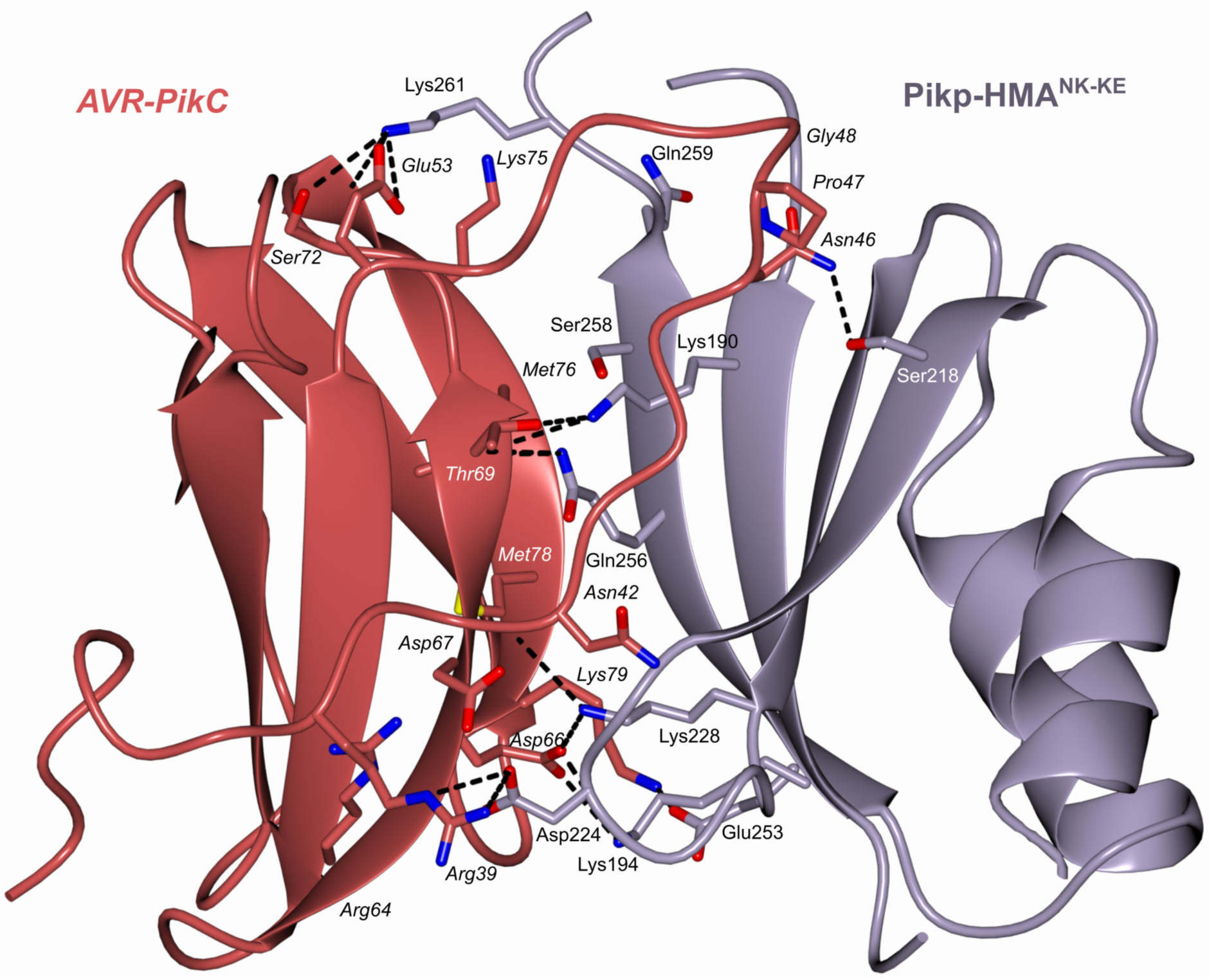
Schematic representation of the crystal structure of the complex formed between Pikp-HMA^NK-KE^ and AVR-PikC (PDB entry 7A8W). The overall structure is similar to other Pik-HMA/AVR-Pik complexes. Amino acid residues forming key contacts at the interface are labelled, including Asp67 that distinguishes AVR-PikC from AVR- PikE.

**Supplementary figure 19.**
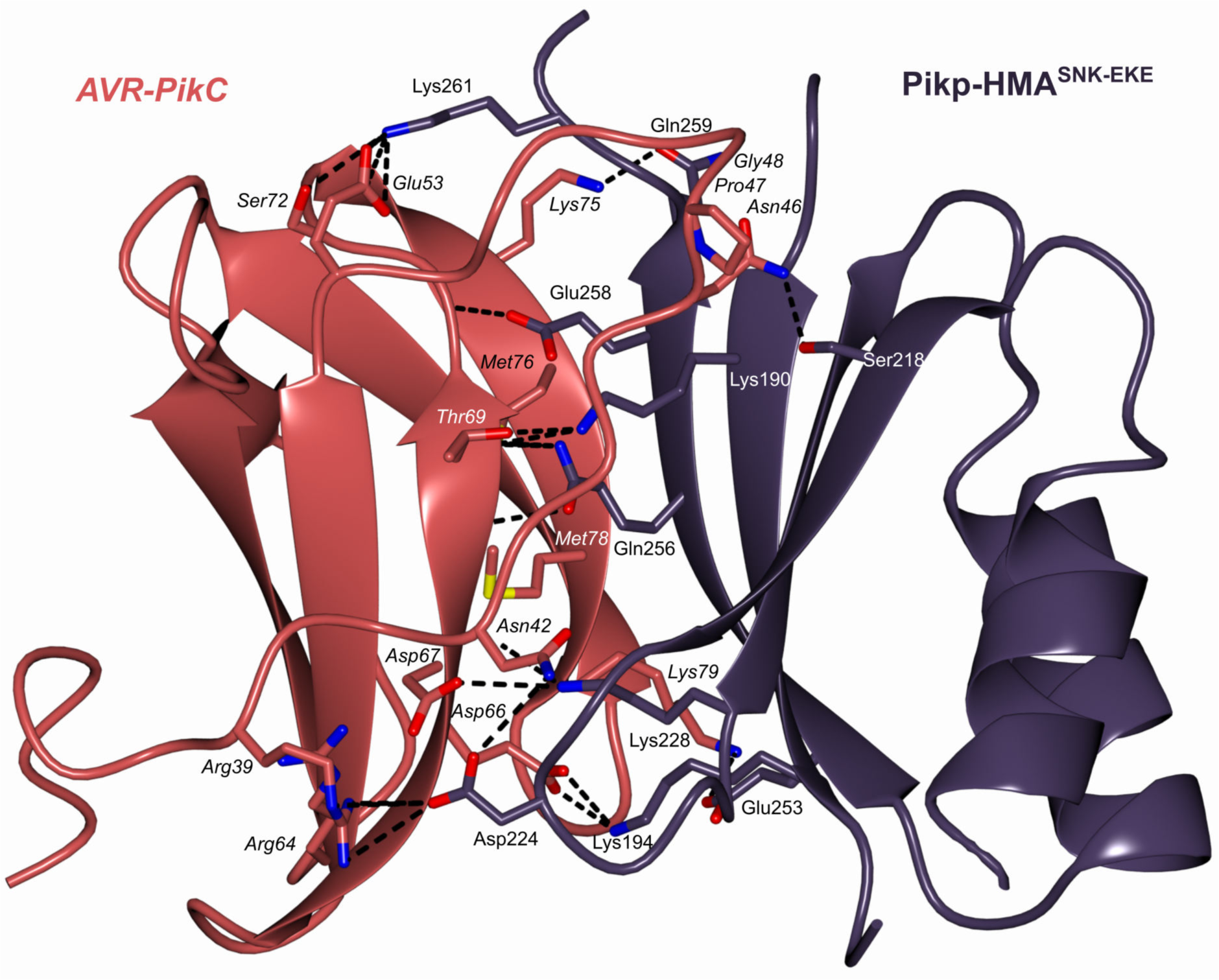
Schematic representation of the crystal structure of the complex formed between Pikp-HMA^SNK-EKE^ and AVR-PikC (PDB entry 7QPX). The overall architecture of the complexes are similar to other Pik-HMA/AVR-Pik structures. Amino acid residues forming key contacts at the interface are labelled, including Asp67 that distinguishes AVR-PikC from AVR-PikE.

**Supplementary figure 20.**
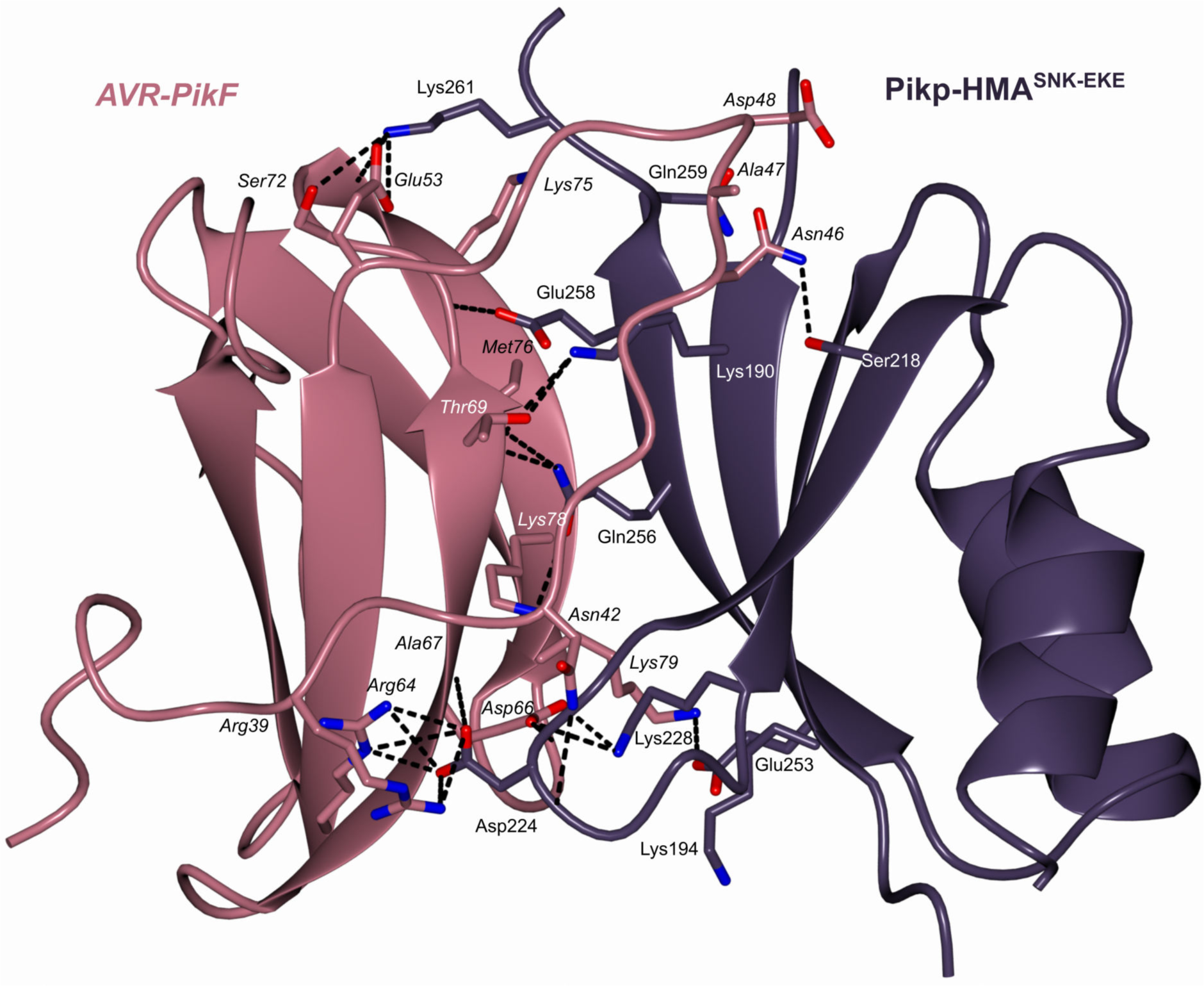
Schematic representation of the crystal structure of the complex formed between Pikp-HMA^SNK-EKE^ and AVR-PikF (PDB entry 7QZD). The overall architecture of the complexes are similar to other Pik-HMA/AVR-Pik structures. Amino acid residues forming key contacts at the interface are labelled, including Lys78 that distinguishes AVR-PikF from AVR-PikA.

**Supplementary figure S21.**
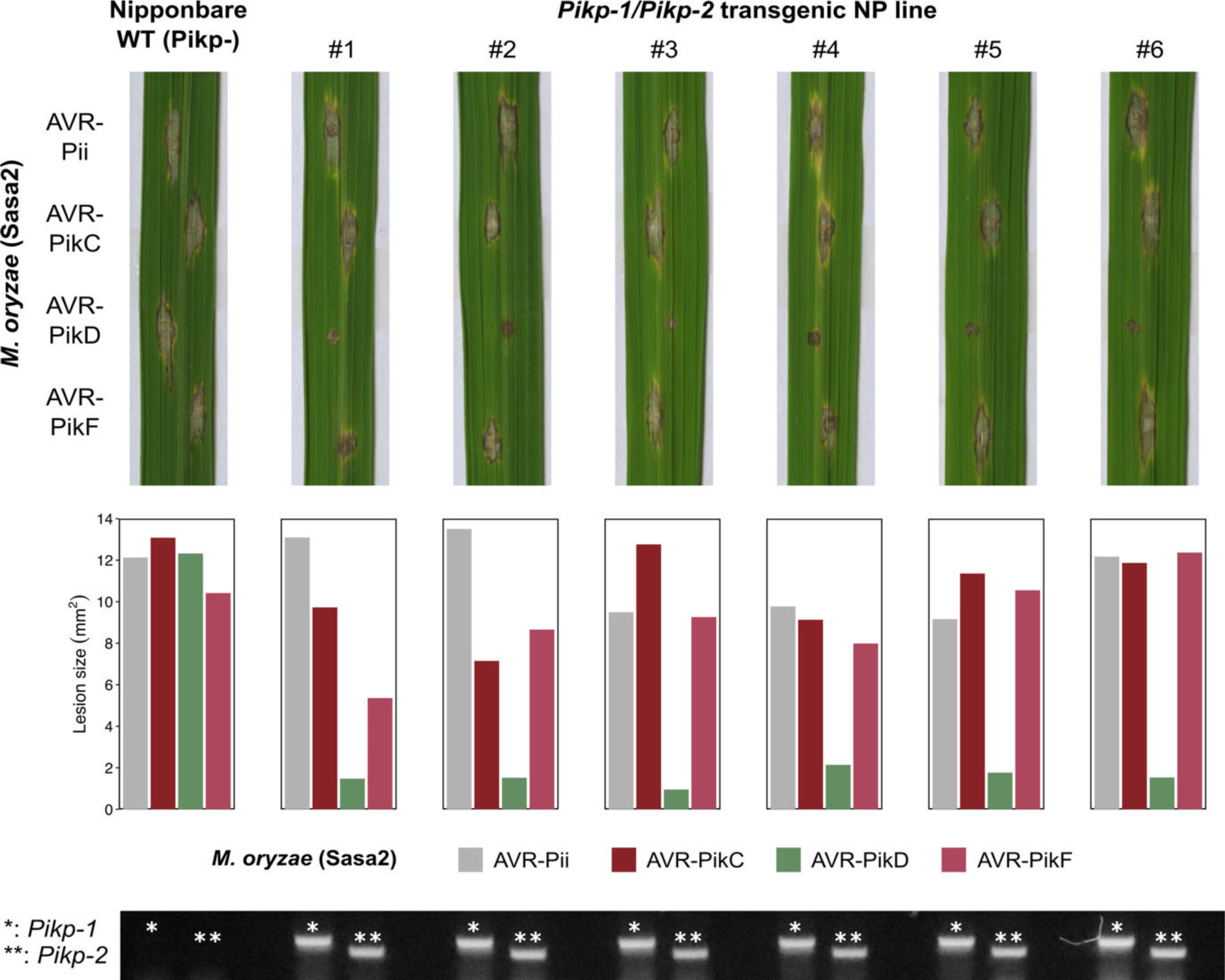
Pathogenicity assays in the T1 progenies derived from a *Pikp- 1/Pikp-2* T0 transgenic line of *O. sativa* cv. Nipponbare against *M. oryzae* Sasa2 transformed with *AVR-Pii*, *AVR-PikC*, AVR*-PikD* or AVR*-PikF*. Images of leaves from different T1 lines were taken 7 days after inoculation. Bar charts show disease lesion sizes (mm^2^) as determined using ImageJ for the specific leaves shown. Gel images show PCR confirmation of transgenes. The plants show susceptibility to all *M. oryzae* strains in the recipient line (WT) and resistance only to AVR-PikD in the transgenics.

**Supplementary figure S22.**
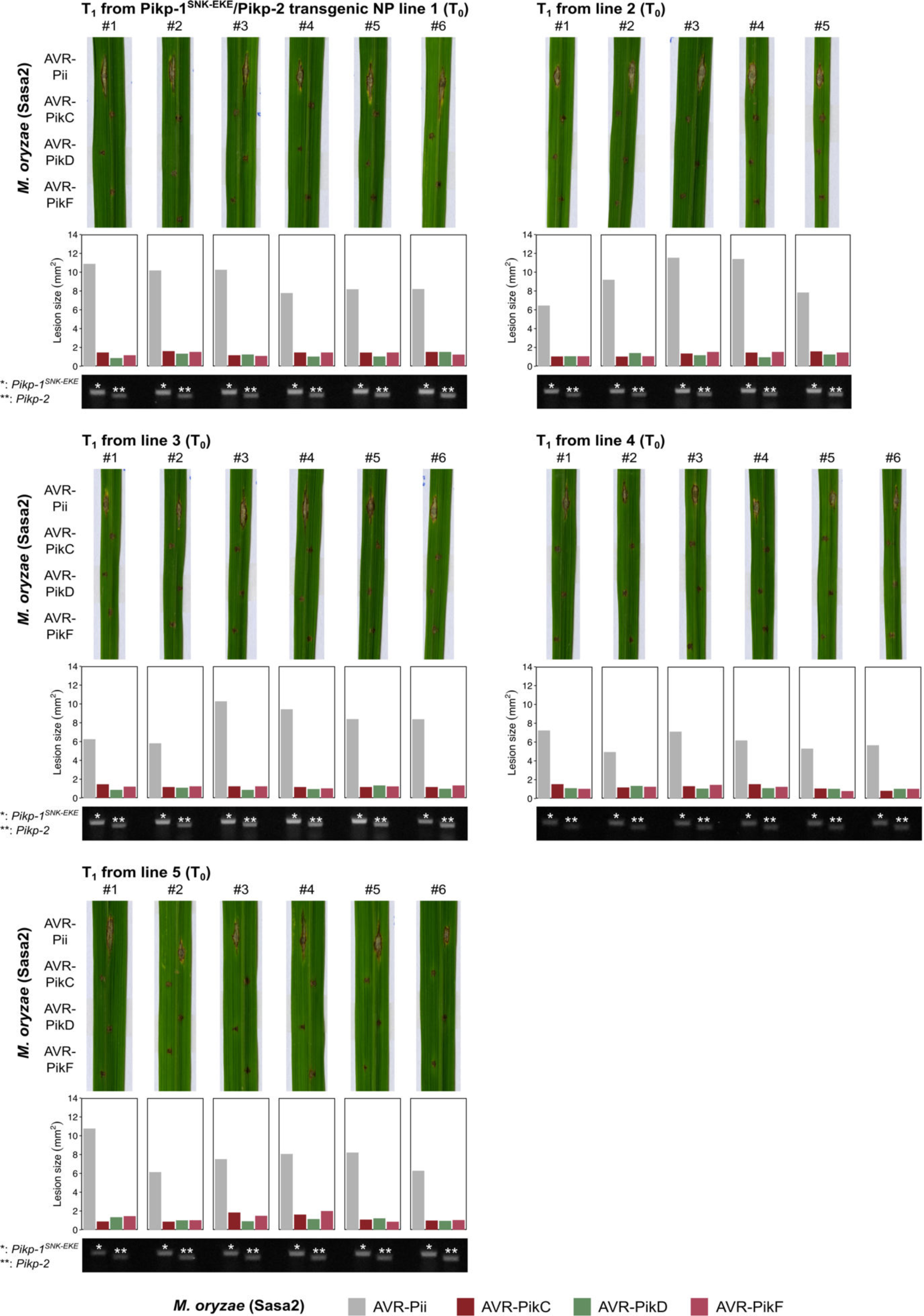
Pathogenicity assays in the T1 progenies derived from five independent *Pikp-1^SNK-EKE^/Pikp-2* T0 transgenic lines of *O. sativa* cv. Nipponbare against *M*. *oryzae* Sasa2 transformed with *AVR-Pii*, *AVR-PikC*, AVR*-PikD* or AVR*-PikF*. Images of leaves from different T1 lines were taken 7 days after inoculation. Bar charts show disease lesion sizes (mm^2^) as determined using ImageJ for the specific leaves shown. Gel images show PCR confirmation of transgenes. The plants show resistance to all *M. oryzae* strains carrying AVR- Pik effectors, but are susceptible to *M. oryzae* Sasa2 carrying AVR-Pii.

**Supplementary figure S23.**
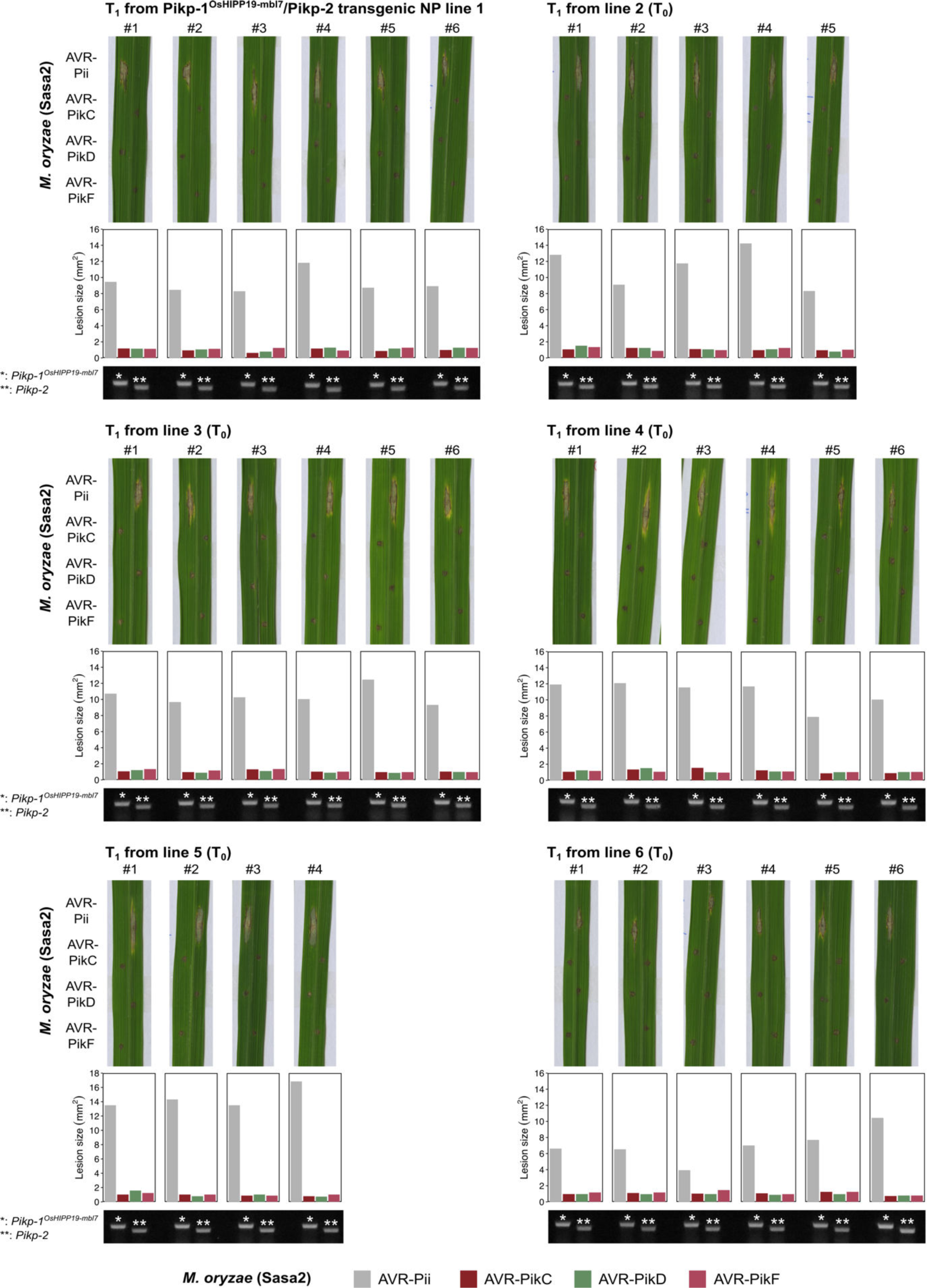
Pathogenicity assays in the T1 progenies derived from six independent *Pikp-1^OsHIPP19-mbl7^/Pikp-2* T0 transgenic lines of *O. sativa* cv. Nipponbare against *M. oryzae* Sasa2 transformed with *AVR-Pii*, *AVR-PikC*, AVR*-PikD* or AVR*-PikF*. Images of leaves from different T1 lines were taken 7 days after inoculation. Bar charts show disease lesion sizes (mm^2^) as determined using ImageJ for the specific leaves shown. Gel images show PCR confirmation of transgenes. The plants show resistance to all *M. oryzae* strains carrying AVR-Pik effectors, but are susceptible to *M. oryzae* Sasa2 carrying AVR-Pii.

**Supplementary figure S24.**
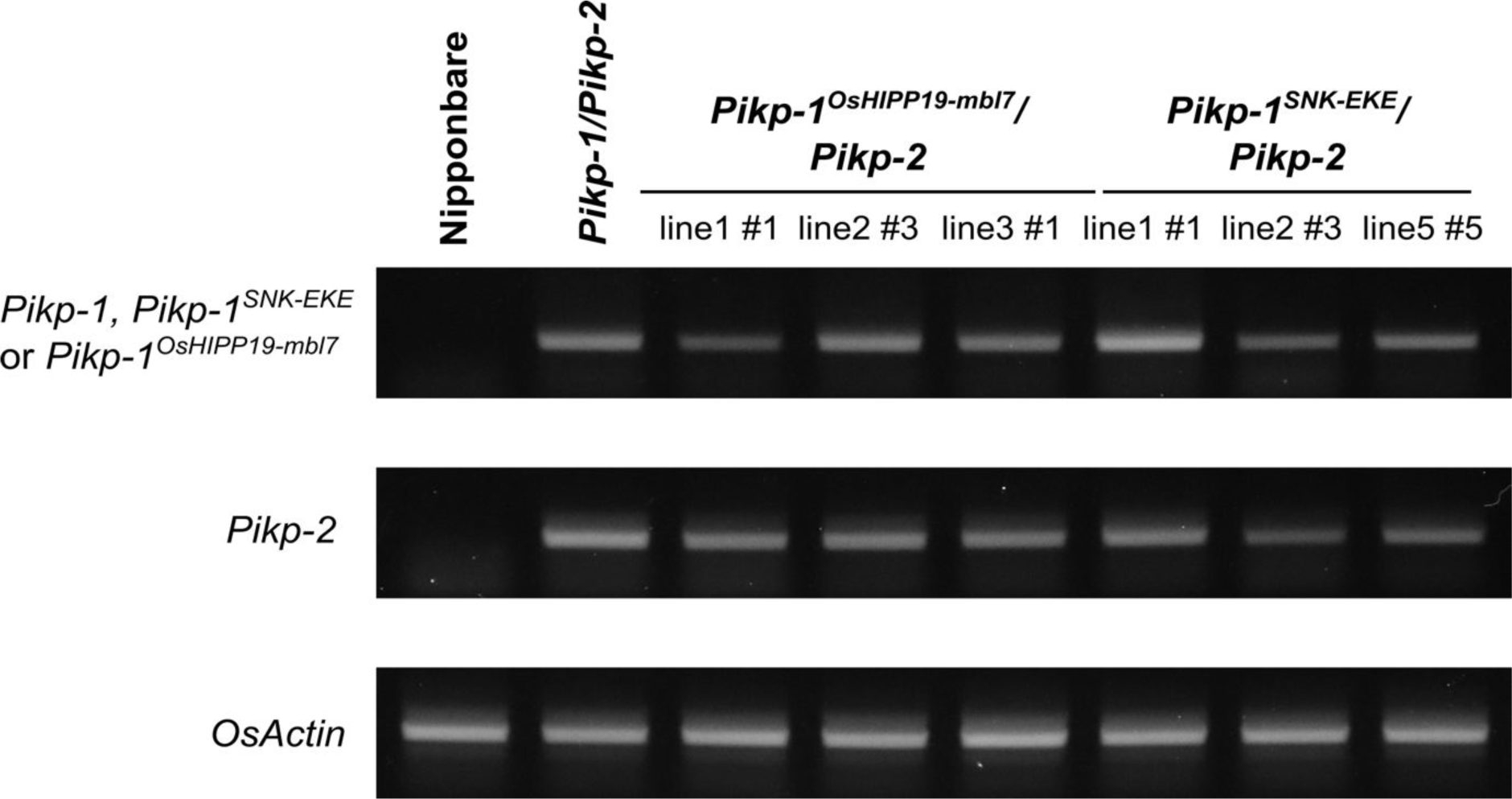
RT-PCR to confirm transgene expression of *Pikp-1* (or engineered *Pikp-1* variants) and *Pikp-2* for the individuals shown in Figure 4.

**Table S1.** X-ray data collection and refinement statistics for Pikp-HMA^NK-KE^/AVR-PikC (PDB entry 7A8W), Pikp-HMA^SNK-EKE^/AVR-PikC (PDB entry 7QPX), and Pikp-HMA^SNK-EKE^/AVR-PikF (PDB entry 7QZD).

|  | Pikp<sup>NK-KE</sup>:AVR-PikC | Pikp<sup>SNK-EKE</sup>:AVR-PikC | Pikp<sup>SNK-EKE</sup>:AVR-PikF |
| --- | --- | --- | --- |
| **Data collection statistics** |  |  |  |
| Wavelength (Å) | 0.9795 | 0.9795 | 0.9795 |
| Space group | *P* 2<sub>1</sub> 2<sub>1</sub> 2<sub>1</sub> | *P* 2<sub>1</sub> 2<sub>1</sub> 2<sub>1</sub> | *P* 2<sub>1</sub> 2<sub>1</sub> 2<sub>1</sub> |
| Cell dimensions: |  |  |  |
| *a*, *b*, *c* (Å) | 66.78, 80.21, 105.68 | 66.35, 83.13, 107.04 | 67.38, 80.78, 103.80 |
| α, β, γ (°) | 90.00, 90.00, 90.00 | 90.00, 90.00, 90.00 | 90.00, 90.00, 90.00 |
| Resolution (Å)* | 46.17-2.15 (2.22-2.15) | 46.67-2.05 (2.11-2.05) | 103.80-2.20 (2.27-2.20) |
| *R*<sub>merge</sub> (%)<sup>#</sup> | 5.3 (99.6) | 8.1 (104.8) | 9.8 (75.1) |
| *I*/*σI*<sup>#</sup> | 23.9 (2.3) | 15.2 (2.0) | 11.5 (2.2) |
| Completeness (%)<sup>#</sup> | 99.9 (99.8) | 99.8 (99.4) | 96.8 (98.6) |
| Unique reflections<sup>#</sup> | 31604 (2694) | 37795 (2886) | 27072 (2013) |
| Redundancy<sup>#</sup> | 13.2 (13.7) | 9.9 (10.1) | 6.9 (6.7) |
| CC<sup>(1/2)</sup> (%)<sup>#</sup> | 100.0 (92.5) | 99.9 (83.8) | 99.9 (84.8) |
| **Refinement/model statistics** |  |  |  |
| Resolution (Å) | 44.16-2.15 (2.21-2.15) | 46.71-2.05 (2.10-2.05) | 63.83-2.20 (2.26-2.20) |
| *R*<sub>work</sub>/*R*<sub>free</sub> (%)<sup>^</sup> | 22.2/27.1 (36.1/34.1) | 20.4/22.7 (27.7/29.5) | 23.1/28.5 (36.3/37.2) |
| No. atoms (Protein) | 6959 | 7016 | 3526 |
| No. atoms (Water) | 78 | 203 | 109 |
| B-factors (Protein) | 64.0 | 46.7 | 44.6 |
| B-factors (Water) | 57.4 | 46.8 | 36.7 |
| R.m.s. deviations:<sup>^</sup> |  |  |  |
| Bond lengths (Å) | 0.008 | 0.008 | 0.009 |
| Bond angles (°) | 1.479 | 1.519 | 1.597 |
| Ramachandran plot (%): <sup>**</sup> |  |  |  |
| Favoured | 97.2 | 98.6 | 97.1 |
| Allowed | 2.8 | 1.4 | 2.7 |
| Outliers | 0.00 | 0.00 | 0.2 |
| MolProbity Score | 1.87 (89<sup>th</sup> percentile) | 1.60 (95<sup>th</sup> percentile) | 1.93 (89<sup>th</sup> percentile) |

\*The highest resolution shell is shown in parenthesis.
<sup>#</sup>As calculated by Aimless, <sup>^</sup>As calculated by Refmac5, <sup>\*\*</sup>As calculated by MolProbity

**Table S2.** Summary of interface analysis by QtPISA for Pikp-HMA^NK-KE^/AVR-PikC (PDB entry 7A8W), Pikp-HMA^SNK-EKE^/AVR-PikC (PDB entry 7QPX), and Pikp-HMA^SNK-EKE^/AVR-PikF (PDB entry 7QZD). Protein chains used for the analysis in each complex (as defined in the PDB entries) are: Pikp^NK-KE^:AVR-PikC (E and F); Pikp^SNK-EKE^:AVR-PikC (E and F); Pikp^SNK-EKE^:AVR- PikF (F and G).

|  |  | Pikp<sup>NK-KE</sup>:AVR-PikC | Pikp<sup>SNK-EKE</sup>:AVR-PikC | Pikp<sup>SNK-EKE</sup>:AVR-PikF |
| --- | --- | --- | --- | --- |
| AVR-Pik | B.S.A. (Å) | 975.4 | 979.2 | 974.1 |
|  | % B.S.A. of total | 18.0 | 18.2 | 18.0 |
| HMA | B.S.A. (Å) | 1019.0 | 1047.0 | 1037.6 |
|  | % B.S.A. of total | 21.5 | 22.4 | 21.4 |
| Total interface area* (Å) |  | 997.2 | 1013.1 | 1005.9 |
| Solvation energy (kcal/mol) |  | -5.2 | -3.2 | -2.4 |
| Binding energy (kcal/mol) |  | -13.6 | -13.2 | -13.6 |
| Hydrophobic p-value |  | 0.4969 | 0.5560 | 0.6137 |
| Hydrogen bonds |  | 12 | 15 | 16 |
| Salt bridges |  | 8 | 9 | 11 |
| Disulphide bonds |  | 0 | 0 | 0 |

\*Total interface area is the total B.S.A. (Buried Surface Area) of each component divided by two.

